# Disruption of theta-timescale spiking impairs learning but spares hippocampal replay

**DOI:** 10.1101/2025.09.15.675587

**Authors:** Abhilasha Joshi, Alison E. Comrie, Samuel Bray, Abhijith Mankili, Jennifer A. Guidera, Rhino Nevers, Xulu Sun, Emily Monroe, Viktor Kharazia, Ryan Ly, Daniela Astudillo Maya, Denisse Morales-Rodriguez, Jai Y. Yu, Anna Kiseleva, Victor Perez, Loren M. Frank

## Abstract

The ability to rapidly learn and retrieve salient information about new environments is critical for survival. In mammals, the hippocampus plays a crucial role in that learning. Specialized features of hippocampal population coding, including network-level theta oscillatory activity, location-specific firing of principal cells, and reactivation of experience during immobility (replay), have been implicated in rapid storage and retrieval of spatial information. Disruptions of theta and replay jointly, or replay alone, are sufficient to impair learning; however, the specific contribution of theta-associated temporal structure during locomotion remains unknown. In this study, we manipulated hippocampal spiking activity in rats specifically during locomotion by optogenetically activating septal parvalbumin-expressing GABAergic neurons. We developed a closed-loop, theta phase-specific stimulation protocol that reliably reduced theta power shortly after stimulation onset. This manipulation preserved the place codes of individual cells but disrupted the fine temporal structure of endogenous spatio-temporal representations (i.e., theta sequences) at the pairwise and population level. Theta disruption during locomotion was also sufficient to cause pronounced deficits in learning the more cognitively challenging component of a spatial alternation task, even though disruption was applied on only *∼*66% of trials. Notably, network effects accompanying theta disruption were restricted to locomotor periods; we did not observe changes in replay rate, length, or content during immobility. Together, these results demonstrate the importance of the precise temporal microstructure of locomotion-associated spatial representations in the hippocampus for learning.

## Introduction

Survival in novel environments is contingent on the acquisition of spatial knowledge and updating behavioral actions based on learned spatial relationships. This learning process engages numerous brain regions and circuits, with the hippocampus playing a critical role in memory-guided spatial navigation, particularly during initial learning in novel environments. Historically, foundational insights into the hippocampus’ role were gained from experiments involving its complete removal or inactivation (Kim and Frank, 2009, Morris et al., 1982, Scoville and Milner, 1957, Kim and Fanselow, 1992). These and many other studies established a critical role for the hippocampus in learning but could not explain why disrupting the hippocampus had such profound effects.

Many efforts to address that question have focused on the population activity patterns thought to mediate hippocampal function. Individual hippocampal pyramidal neurons are active in specific locations (’place cells’ firing in ‘place fields’) (O’Keefe, 1976). Place firing during movement is also closely associated with *∼*8 Hz theta oscillations: pyramidal cells tend to fire most during the descending phases of theta and least during the ascending phases (Mizuseki et al., 2009). Furthermore, at a population level, each cycle of the theta rhythm typically engages a sequence of place cell firing (a ‘theta sequence’) (Skaggs et al., 1996, Foster and Wilson, 2007, Burgess et al., 1994). This sequence relates theta to the content of spatial representations, such that positions just behind and at the current position of the animal are represented along the descending phases of theta, while alternative “non-local” positions, such as those further ahead, are regularly expressed during ascending phases (Foster and Wilson, 2007, Kay et al., 2020, Joshi et al., 2023, Gupta et al., 2012).

Subsequently, during periods of awake immobility and sleep, the hippocampus frequently reactivates patterns associated with awake experience (Diba and Buzsáki, 2007, Joo and Frank, 2018, Buzsáki et al., 1983, Panoz-Brown et al., 2018, Karlsson and Frank, 2009). Such reactivation events, which often occur during a bout of high frequency oscillatory activity called sharp-wave ripples (SWRs), can include sequential replay of multiple place cells that together represent coherent spatial trajectories through an environment (Diba and Buzsáki, 2007, Pfeiffer and Foster, 2013, Kay et al., 2016, Karlsson and Frank, 2009, Davidson et al., 2009).

Theta- and SWR-related sequences are well suited to contribute to learning and memory-guided spatial navigation, both through the potential induction of plasticity in downstream structures and through the activation of spatially remote representations that could support flexible decision-making (Buzsáki, 1989, Hasselmo, 2025, Skaggs et al., 1996, Foster and Wilson, 2007, Jadhav et al., 2012, Gillespie et al., 2021, Denovellis et al., 2021, Johnson and Redish, 2007, Kay et al., 2020, Pastalkova et al., 2008, Comrie et al., 2024). At the same time, the precise role of these sequences in learning and decision-making remains unclear.

Previous work has established that manipulations targeting local field potential (LFP) patterns correlated to these sequences can disrupt navigational behaviors without substantively affecting overall spatial tuning. Studies related to theta have used manipulations of neurons in the medial septum (MS), a brain region critical for generating and regulating the hippocampal theta rhythm, and for spatial learning (Mizumori et al., 1990, Robinson et al., 2023). Specifically, disruptions of normal theta oscillatory activity, either through pharmacological inactivation using muscimol (Wang et al., 2015, 2016), cooling of the MS and reducing theta frequency (Petersen and Buzsáki, 2020), entrainment of the endogenous LFP during movement at a frequency well above the normal *∼*8 Hz (Zutshi et al., 2018, Quirk et al., 2021), or scrambling of theta (Etter et al., 2023), impair performance in a previously learned task. These manipulations typically preserve place field tuning and size, (Koenig et al., 2011, Brandon et al., 2014, Robbe and Buzsáki, 2009) (but see (Wang et al., 2015)), although overall firing rates are reduced (Koenig et al., 2011, Brandon et al., 2014, Drieu et al., 2018).

Disrupting SWR activity or elongating SWR duration during awake immobility also affects behavior, impairing or improving learning in a spatial alternation task, but does not substantially alter place fields. (Jadhav et al., 2012, Fernández-Ruiz et al., 2019). Effects on theta sequences were not reported in these studies, but these results are broadly consistent with a role for replay and reactivation in learning. In a key recent study, optogenetic activation of medial entorhinal inhibitory neurons at gamma frequencies during locomotion was shown to disrupt theta sequences, subsequent sequential replay, and the ability of rats to learn to navigate efficiently toward rewarded locations in a maze (Liu et al., 2023). Another study showed that a behavioral manipulation (e.g. passive movement in a trolley) disrupts theta oscillations and associated spatial sequences and SWR-associated replay (Drieu et al., 2018). Both these studies suggested that theta sequences are necessary to establish replay sequences, raising the possibility that disruptions of theta may affect learning through an impact on replay.

These findings contrasted with earlier work, which used muscimol inactivation of the MS and showed that disruption of theta oscillations and spatial sequences during locomotion did not abolish sharp wave ripples and associated replay, suggesting that a dissociation between the two was possible (Wang et al., 2015, 2016). In this study, the ability of animals to perform a previously learned task was also impaired, and the authors suggested that the importance of replay sequences might be limited to learning a new environment or a new aspect of a task. That said, the rate of ripple events in their study increased during the muscimol inactivation, raising the possibility that an increase in the number of replays may also contribute to impairment in learning by providing conflicting information to downstream brain areas.

Taken together, these results demonstrate the importance of both theta sequences and replay sequences, but leave open the question of whether these two phenomena can be dissociated in terms of their impact on learning. Specifically, we do not know whether theta oscillatory activity and theta sequences can be selectively disrupted at short timescales, without affecting spatial tuning and replay, and whether such a targeted disruption alone would be sufficient to impair learning? Answering this question requires a manipulation that is both more selective and more temporally precise than those previously attempted, and an experimental and analytical approach that could characterize theta dynamics, place cell properties and replay in the same animals and sessions.

Addressing this question is of particular interest because hippocampal activity in the theta state, during locomotion, has been hypothesized to directly support plasticity for learning and memory (Buzsáki, 1989, Skaggs et al., 1996, Jensen and Lisman, 1996, Wang et al., 2015, Mizuseki et al., 2009, O’Keefe and Recce, 1993, Hasselmo et al., 2002, Comrie et al., 2022, Foster and Wilson, 2007). Additionally, understanding how individual physiological features, such as these locomotion-associated theta dynamics, contribute to learning may also enable identification of therapeutic strategies for improving memory in the setting of hippocampal dysfunction (Gillespie et al., 2024, Atherton et al., 2015, Fernández-Ruiz et al., 2019, Martorell et al., 2019). We therefore developed a closed-loop approach to reliably manipulate hippocampal theta and associated activity during locomotion. Our approach leveraged a specific MS parvalbumin-expressing GABAergic projection, which in turn targets GABAergic neurons in the hippocampal formation (Zutshi et al., 2018, Quirk et al., 2021, Joshi et al., 2017, Wang et al., 2015, Fuhrmann et al., 2015, Robinson et al., 2016, Kloc et al., 2020, Mouchati et al., 2020, Lepperød et al., 2021, Etter et al., 2023, Király et al., 2023, Gemzik et al., 2021, Yong et al., 2022, Robinson et al., 2024). Using this approach, we were able to selectively suppress theta power and disrupt the sequential firing of place cells without affecting average place field activity. This manipulation, when applied in *∼*2/3rds of trials, was sufficient to profoundly impair learning when choices depended on memory for specific past experiences. Learning of simpler spatial trajectories remained intact, as did the expression of replay sequences. These results establish that manipulation of theta oscillatory activity and the associated temporal order of spiking is sufficient to impair learning.

## Results

Medial septal (MS) parvalbumin (PV)-expressing neurons coordinate theta rhythmic activity across the hippocampal formation (Petsche et al., 1962, Freund and Antal, 1988, Viney et al., 2018, Joshi et al., 2017). Previous optogenetic manipulations in mice (Bender et al., 2015, Zutshi et al., 2018, Quirk et al., 2021, Etter et al., 2023) have used optogenetic manipulations of this circuit to entrain hippocampal local field potential at experimenter-requested frequencies between 4 and 20 Hz. We similarly used a viral strategy to express a light-activated channel (ChR2) in PV+ neurons within the MS of PV-Cre transgenic rats. We also implanted an optic fiber targeting the MS as well as either silicon electrodes or multi-tetrode arrays targeting the hippocampus in *n* = 10 rats.

Each of these animals received optical stimulation restricted to periods of locomotion on either a linear track, a W-shaped track, or both (**see methods**). The stimulation pattern was governed by a protocol that either activated MS PV neurons at a wide range of requested frequencies (“rhythmic stimulation”) or at specific phases of the ongoing theta rhythm (“phase-specific stimulation”) recorded in the hippocampus (**Figure 1A**). To enable us to assess whether the effects of the manipulation were restricted to stimulation intervals, each behavioral epoch was divided into three approximately equal periods, an initial and final stimulation interval (henceforth *“stimulation-on”*) and a middle no-stimulation control interval (henceforth *“stimulation-off”*), during which animals ran for rewards without optical stimulation (**Figure 1B**).

**Figure 1.**
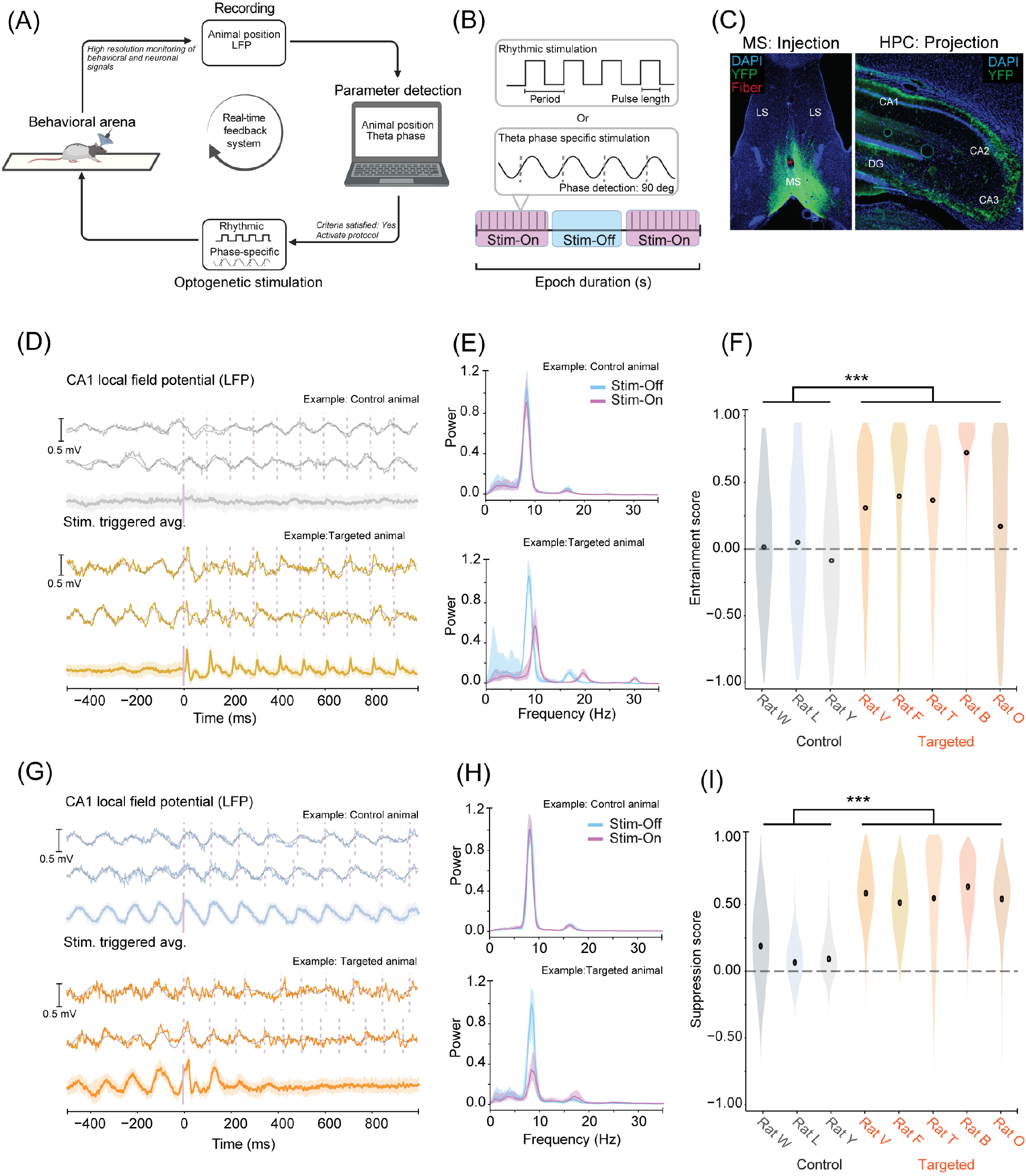
Reliable and reversible manipulation of hippocampal theta oscillations on a linear track. **A**, Schematic of experimental approach. Rats implanted with electrodes targeted to hippocampal region CA1 and a tapered optic fiber targeted to the dorso-ventral extent of the medial septum. Optogenetic stimulation was applied only when the rats were locomoting on a behavioral arena (a linear track shown), using a real-time closed-loop system. **B**, Two types of optogenetic stimulation protocols, either rhythmic stimulation (defined by period and pulse length), or a theta phase-specific stimulation (90-degree targeting shown), was applied in each session (upper grey boxes). A day consisted of interleaved run and rest epochs of 15–20 minutes each. Run epochs included ‘test’ intervals (purple, “*stimulation-on*”), where the optogenetic stimulation would be applied during locomotion, and “control” intervals (blue, “*stimulation-off*”), where no optogenetic manipulation was applied during locomotion. **C**, Virus (AAV5 Ef1a DIO-hChR2(H134R), labeled with eYFP) targeted the medial septum (left) and long-range projecting axon fibers in the hippocampus (right). **D**, Individual examples (upper two) and epoch-averaged (lower) hippocampal CA1 LFP traces during 10 Hz rhythmic stimulation in one control animal (grey) and one targeted animal (orange) triggered by stimulation onset. Error bars show 95% confidence intervals (CI). Smooth traces overlaying raw LFPs are band-pass filtered between 5 and 11 Hz. **E**, Power spectral density of hippocampal LFP during 10 Hz stimulations during stimulation-on and stimulation-off intervals in one control animal (upper) and one targeted animal (lower). **F**, Entrainment scores distributions across sessions for *n* = 3 control animals (grey) and *n* = 5 targeted animals (orange). Targeted animals show significant entrainment compared to controls (mixed effect linear model (animal, transfection state), effect of transfection; *p* = 0.0015, targeted *n* = 19816 interval pairs, control *n* = 8517 interval pairs). **G**, Individual examples (upper two) and epoch-averaged (lower) hippocampal CA1 LFP traces during theta phase-specific stimulation in one control animal (grey) and one targeted animal (orange) over time since stimulation onset. Error bars show 95% CI. Smooth traces overlaying raw LFPs are band-pass filtered between 5 and 11 Hz. **H**, Power spectral density of hippocampal LFP during theta-phase specific manipulation during stimulation-on and stimulation-off intervals in one control animal (upper) and one targeted animal (lower). **I**, Suppression score distributions across sessions for *n* = 3 control animals (grey) and *n* = 5 targeted animals (orange). Targeted animals show significant suppression compared to controls (mixed effect LM (animal, transfection state), effect of transfection *p* = 2.6 *×* 10*^−^*^27^, targeted *n* = 34268 intervals, control *n* = 14250 intervals). Note, one control animal (Rat I) lacked sufficient runs, and one targeted animal (Rat D) was not tested at the highest laser power during locomotion. Hence these animals could not be included in the entrainment and suppression score calculations.

We confirmed that the PV cells transfected in the MS projected to the hippocampus as expected (**Figure 1C**). Histological analysis also revealed accurate targeting of both virus and optic fiber to the MS in n=6 animals (hereafter the *“targeted”* group) and inaccurate targeting of either virus or optic fiber in n=4 animals (hereafter the *“control”* group). Thus, with this experimental approach, for each experiment, we had both within-animal controls (*stimulation-on* versus *stimulation-off* intervals) and across-animal controls (*targeted* versus *control*). In a subset of animals (n=3), we also determined the laser power and pulse durations required to reliably entrain hippocampal LFP during locomotion and rest (**see Methods**). For each analysis, we included all animals in which the relevant manipulation was tested and that met the predefined inclusion criteria (**see Methods**).

### Reliable and reversible manipulation of hippocampal theta oscillations on linear track

In targeted but not control animals, rhythmic optogenetic stimulation of MS PV neurons reliably entrained hippocampal LFP oscillations during *stimulation-on* intervals without any detectable effect on endogenous theta activity during *stimulation-off* intervals (**Figure 1D**, example of 10 Hz stimulation). Correspondingly, the power spectrum of the LFP during *stimulation-on* intervals (**Figure 1E**) showed a marked peak at the targeted frequency. To quantify the degree of entrainment, we computed an *entrainment score*, which was defined as the excess LFP power observed at the requested frequency during stimulation periods compared to control periods (**see Methods**). We found a robust entrainment of the LFP at requested frequencies for targeted animals and no detectable change in control animals (**Figure 1F**, example of 10 Hz). These observations were consistent across stimulation frequencies (**Supplementary Figure 1**). These experiments replicate results from previous work in transgenic mice (Zutshi et al., 2018, Quirk et al., 2021, Etter et al., 2023) and helped us optimize the optic fiber, laser power, and pulse widths required for effectively driving MS neurons in transgenic rats (**see Methods**).

### Closed-loop theta phase-specific manipulation results in a rapid reduction of theta power on linear track

Spiking activity in the hippocampus is closely associated with ongoing theta oscillations, with pyramidal cells tending to fire least during the ascending phases of theta and most during the descending phases (Mizuseki et al., 2009). This temporal structure is coordinated by a network of local and long-range GABAergic inputs from cortical and subcortical regions (Joshi et al., 2017, Varga et al., 2012, Viney et al., 2018, Salib et al., 2019, Király et al., 2023), each precisely timed and phase-coupled to endogenous theta oscillatory activity. A subset of rhythmic MS PV neurons, in particular, project to hippocampal axo-axonic cells (Viney et al., 2013, Joshi et al., 2017), and thus activation of MS PV neurons should have a disinhibitory effect on hippocampal pyramidal cells. We therefore reasoned that closed-loop activation of MS PV neurons during the ascending phase of ongoing theta (**Supplementary Figure 2A**) would introduce additional pyramidal cell spikes and disrupt normal hippocampal theta dynamics.

Consistent with this prediction, we found optogenetically activating MS PV neurons at the ascending phase of theta resulted in a marked reduction of theta power beginning approximately 200 ms after initial stimulation onset (**Figure 1G**). To quantify this relationship, we computed a *suppression score*, defined as the decrease in LFP power observed at the peak frequency during *stimulation-off* periods compared to that observed during *stimulation-on* periods (**see methods**). This measure confirmed the prominent reduction in theta power (**Figure 1H**) in targeted animals compared to control animals (**Figure 1I**). As in the rhythmic stimulation condition, these changes were restricted to stimulation times, and endogenous theta dynamics were unaffected during *stimulation-off* run periods.

This manipulation did not affect running velocity between *stimulation-on* intervals compared to *stimulation-off* intervals (*t*-test between the distribution of average velocities per-trial within each animal group; targeted animals: *p* =0.8; control animals: *p* =0.5). Thus, this theta-phase-specific manipulation selectively suppresses the extracellularly measured theta rhythm while leaving gross locomotor behavior intact.

### Theta disruption alters temporal spiking of pyramidal cells, but preserves place fields

The disruption of theta oscillations could potentially affect both temporal and spatial properties of individual hippocampal pyramidal cells. Previous studies have yielded varied results, with some dissociating spatial and temporal properties through specific optogenetic manipulations (Liu et al., 2023), and others reporting mixed effects (Etter et al., 2023). To examine the effects on temporal and spatial coding we first verified that putative pyramidal cells in control animals were unaffected by the light pulses while cells in targeted animals showed short latency (5–10 ms), robust responses (**Figure 2A-C**). This effect is consistent with the strong disinhibitory input from the medial septum to the hippocampus (Joshi et al., 2017) and the expected synaptic delays by strongly myelinated GABAergic neurons (Jones et al., 1999). Note, that while hippocampal cells respond within 5-10ms of the manipulation, theta power appeared to decrease gradually over a period of hundreds of milliseconds. This difference is consistent with the view that theta oscillatory activity that can be measured in the hippocampus is a result of a multi-region feedback loop that involves various cortical and subcortical networks, a feature that may make it robust to individual perturbations.

**Figure 2.**
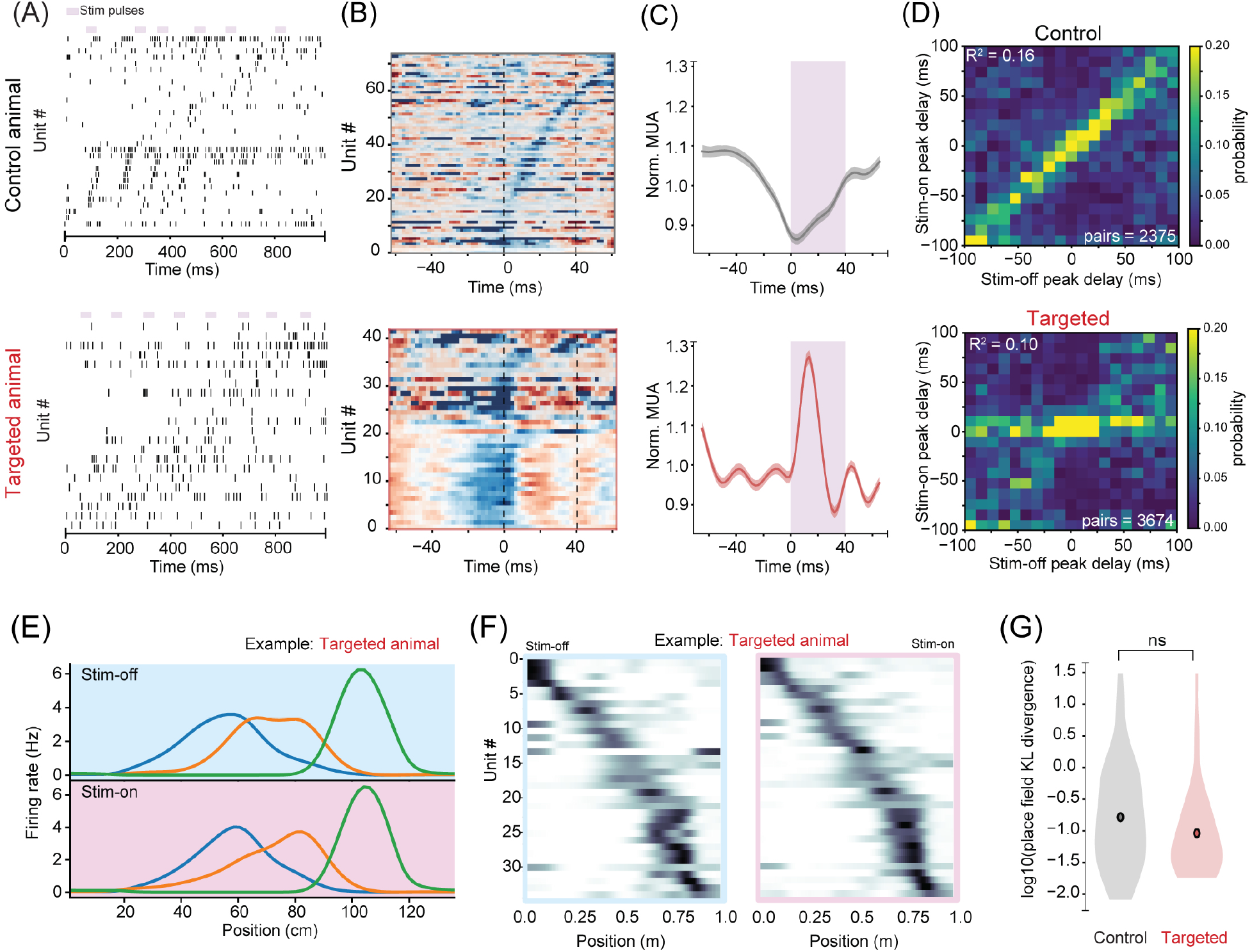
Theta disruption impairs temporal coding of pyramidal cells, but preserves place code on linear track. **A**, Raster plots of simultaneously recorded hippocampal neurons ordered by the peak of their place fields with MS optogenetic theta phase-specific stimulation pulses highlighted (purple) in one control (top) and one targeted (bottom) animal. Note that in control rats, endogenous theta sequences are apparent. **B**, Stimulation-triggered averages of simultaneously recorded hippocampal neurons show the absence of consistent stimulation-triggered organization in control animal (top) and entrainment of spiking by optogenetic activation in targeted animal (bottom). Vertical dashed lines indicate stimulation time. Firing rates (normalized) shown in heatmap. **C**, Stimulation-triggered MUA across all control (*n* = 499 units, 4815 pulses, top) and targeted (*n* = 583 units, 5143 pulses, bottom) animals shows distinct MUA spiking structure during stimulation. In control animals, the expected relationship is observed, with spiking probability decreasing during stimulation times that occur in the ascending theta phases, whereas in targeted animals spiking probability increases *∼*10 ms after stimulation onset. Shaded regions are 95% CI of the mean. **D**, Pairwise temporal offsets between cross-correlograms computed in *stimulation-on* versus *stimulation-off* intervals in control (*n* = 3674 pairs, top) and targeted (*n* = 2375 pairs, bottom). Note, a strong diagonal indicates that the cross-correlation structure is maintained in control animals (as expected). Control vs. targeted *p <* 10*^−^*^4^ for pooled data; hierarchical bootstrap *p* = 0.15. **E**, Examples of three place cells’ spatial selectivity in targeted animals during *stimulation-off* (blue) versus *stimulation-on* (purple) intervals. Note similar spatial tuning patterns. **F**, Spatial stability across all identified place cells in one targeted animal during *stimulation-off* (left, blue box) and *stimulation-on* (right, purple box) periods. Note similar spatial tuning patterns. **G**, Place field stability is measured for all units as the KL divergence (**see Methods**) of their spatial firing selectivity between *stimulation-on* and *stimulation-off* intervals (*t*-test of distributions of KL divergence between targeted (*n* = 35 units satisfying inclusion criteria across *n* = 3 animals) and control (*n* = 43 satisfying inclusion criteria across *n* = 4 animals) animals, *p* = 0.8).

Our manipulation disrupted theta, which meant that we could not carry out standard analyses that quantify the relationship between theta phase and spiking. We therefore took a different approach where we examined the sequential temporal relationships between principal cells that is typically present in conjunction with theta modulation. To do so we constructed cross-correlation histograms for pairs of putative place neurons (i.e., units with localized place fields; **see Methods**) during both *stimulation-on* and *stimulation-off* intervals. These histograms typically exhibit strong theta modulation, with peaks at the theta timescale; the offset of these peaks from zero reflects longer timescale differences in firing related to spatial firing fields (Dragoi and Buzsáki, 2006, Skaggs et al., 1996). For each cross-correlation, we identified the highest peak value within a ±100 millisecond lag. We then pooled data across all animals and run sessions and plotted the location of each cell pair’s peak during *stimulation-on* versus *stimulation-off* intervals (**Figure 2D**).

These plots revealed the expected diagonal of peak offset for control animals, reflecting consistent cross-correlation structure with or without stimulation. Consistent structure was less readily visible in targeted animals, with many cell pairs showing more synchronous activity during *stimulation-on* periods (*R*^2^ = 0.10, control vs. targeted p < 10*^−^*^4^ for pooled data; more conservative hierarchical bootstrap p = 0.15, reflecting low sample count for some animals).

By contrast, we did not detect any changes in the spatial tuning of putative pyramidal neurons. Specifically, the increased spiking during *stimulation-on* intervals respected place field boundaries (**Supplementary Figure 2B**) and the place fields of single cells were not detectably affected by theta disruption (**Figure 2E-F**, single cell place fields, pooled for **G**). To assess these observations at the population level, we computed the similarity between the place field peaks in control versus targeted animals. We could not detect a difference between the two conditions (Spearman correlation of place field peaks between stim-on and stim-off intervals: p= 0.68, t-test). These results are consistent with previous studies that performed MS manipulations (Zutshi et al., 2018, Etter et al., 2023). Thus, our manipulation protocol provides an opportunity to ask specifically how temporal coding contributes to behavior.

### Theta disruption impairs learning in a novel spatial environment

We next investigated the effects of disruption of locomotion-associated hippocampal temporal dynamics on learning. Specifically, we focus on the initial learning of a hippocampal-dependent spatial alternation task in a W-track environment (Kim and Frank, 2009). As in previous experiments on the linear track, we maintained the *stimulation-on*, *stimulation-off*, *stimulation-on* structure (**Figure 1B**) to assess the impact of stimulation on single-cell and population-level activity. In contrast to the linear track experiments, theta-disruption was begun as soon as animals were first put on the maze, allowing us to assess the importance of theta-timescale structure for learning. Here the control group was expanded to include data from our prior work (Joshi et al., 2023) that used the same training protocol.

Despite the interleaved *stimulation-off* periods, targeted animals had pronounced deficits in learning the spatial alternation task. We first examined the more cognitively challenging outbound trials in which animals must remember their previous outer arm visit to make the next correct choice. While control animals readily learned to alternate correctly, targeted animals were profoundly impaired (**Figure 3A, 3B**, *t*-test performance accuracy, p=0.01). By contrast, we did not observe any significant differences in learning across groups on the interleaved inbound trials, where a simpler place–action association is sufficient to enable learning (**Figure 3C, 3D**, *t*-test, performance accuracy, *p* =0.126).

**Figure 3.**
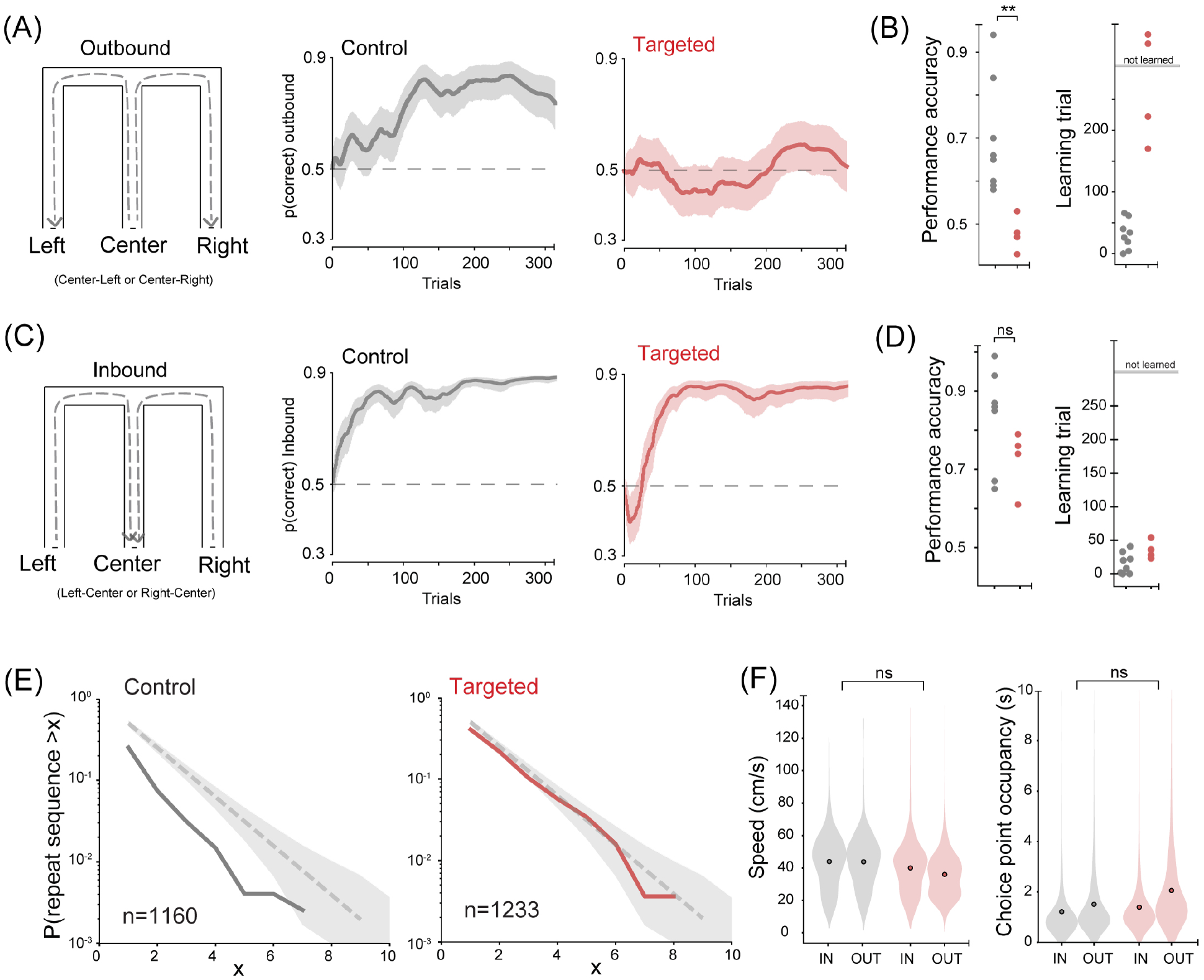
Theta disruption impairs learning of outbound trial structure in a novel W-track environment. **A**, W-track schematic for Outbound trials (left). Learning curve for control animals (*n* = 4, middle, grey) and targeted animals (*n* = 4, right, red) on Outbound trials. **B**, Performance accuracy (left) and learning trial (right) for Outbound trials (*n* = 8 control animals, grey; *n* = 4 targeted animals, red; 4 control animals from this study, 4 control animals from Joshi et al. (2023)). Performance accuracy *t*-test *p* = 0.010. **C**, W-track schematic for Inbound trials (left). Learning curve for control animals (*n* = 4, middle, grey) and targeted animals (*n* = 4, right, red) on Inbound trials. **D**, Performance accuracy (left) and learning trial (right) for Inbound trials (*n* = 8 control animals, grey; *n* = 4 targeted animals, red; *n* = 4 control animals from this study; *n* = 4 control animals from Joshi et al. (2023)). Performance accuracy *p* = 0.126. **E**, Probability of “repeat” error trials (re-visiting a reward port on outbound trials) in control animals (grey solid line, left, *n* = 1160 trials) and targeted animals (right, red solid line, *n* = 1233 trials) compared to the 95% CI (grey shaded region) of an analytical chance distribution. Targeted animals’ choices are not different from the chance distribution. **F**, *Left:* Distribution of running speeds in control (grey) and targeted (red) animals on inbound and outbound trials (linear mixed effects model: effect of targeting, *p* = 0.61; interaction (targeting *×* trial type), *p* = 6.52 *×* 10*^−^*^4^) during time spent outside reward ports. *Right:* Distribution of occupancy times around the choice point in control (grey) and targeted (red) animals on inbound and outbound trials (linear mixed effects model: effect of targeting, *p* = 0.94; interaction (targeting *×* trial type), *p* = 1.39 *×* 10*^−^*^4^).

To further characterize the selective deficit on outbound trials we calculated the probability that animals revisited the same reward port on consecutive trials (“repeats,” **Figure 3E**), which reflects a violation of the task rules and never results in reward. Control animals showed a strong bias to avoid visits to the immediately previously visited arm and thus had a distribution of repeated visits very different than random. By contrast, the targeted animals’ outbound choices were not distinguishable from random, and thus showed no overall history dependence.

This learning disruption on the W-track could not be explained by consistent changes in motivation: both targeted and control animals completed >300 trials, sufficient to measure learning (Jadhav et al., 2012, Joshi et al., 2023). There were similarly no significant differences in average running speed or choice point occupancy during locomotion, but targeted animals did move more slowly and spend more time near the choice point on outbound trials (**Figure 3F, Video 1**, Speed targeted: median: 38.5 cm/s (IQR 25.8–47.1); control: median 46.44 cm/s (IQR 32.8–55.1), linear mixed effects model: effect of transfection; *p* =0.68; interaction (transfection x trial type; p<0.05); occupancy targeted: median: 1.5 s (IQR 1–2.5), control: 0.7 s (IQR 1–1.7), linear mixed effects model: effect of transfection: *p* =0.94, interaction (transfection x trial type; p<0.05)). This pattern is consistent with the targeted animals having greater choice uncertainty on outbound trials.

### Disruption of theta sequences in the W-track

We then asked how our manipulation affected the structure of the spiking activity on the W-track. We first verified that, like on the linear track (**Figure 2C**), stimulation drove an increase in spiking in the targeted, but not the control group (**Supplementary Figure 3A**). We also asked whether there was place field remapping between *stimulation-off* and *stimulation-on* conditions. As on the linear track, we found no difference between place field peak locations (Spearman correlation of place field peaks between stim-on and stim-off intervals: p=0.91, t-test, n=36 control, n=18 targeted epochs).

We next examined the consistency of pairwise cross-correlation peaks between *stimulation-on* and *stimulation-off* periods as we did for the linear track (**Figure 2D**). Here again we observed more consistent cross-correlation structure in control animals (linear fit R^2^ = 0.09), and less consistent structure in targeted animals ( R^2^ = 0.005; **Supplementary Figure 3B**). These differences were highly significant when data were pooled across animals and run epochs (control vs. targeted p < 10*^−^*^4^) and were also significant with the more conservative hierarchical bootstrap (p < 10*^−^*^3^; **Supplementary Figure 3C**).

The presence of a persistent learning deficit on outbound trials despite *stimulation-off* periods suggested the possibility that the suppression of theta in the W-track led to persistent deficits in the temporal structure of hippocampal activity. We therefore compared the structure of activity across control and targeted animals for both *stimulation-on* and *stimulation-off* periods. We plotted, for each cell pair, the peak of the theta time-scale cross-correlation on the y-axis and the peak of the behavioral (*∼*1 second) time-scale cross-correlation on the x-axis (**see Methods**).

These plots revealed a persistent deficit in the temporal organization of spiking in targeted animals present in both *stimulation-on* and *stimulation-off* periods (**Supplementary Figure 3D**). In control animals, these plots showed the expected correlation between short- and long-timescale peaks consistent with theta sequences ( R^2^ = 0.077) but this correlation was largely absent in the targeted animals ( R^2^ = 0.007; pooled comparison p < 10*^−^*^4^, hierarchical bootstrap p < 0.05, **Supplementary Figure 3E**). Thus, disruption of theta during stimulation periods was sufficient to disrupt sequential spiking activity even after stimulation was turned off.

Finally, we also verified that, as in the linear track (**Figure 2G)**, this disruption was confined to fine timescale activity: place fields on W-track remained similar across *stimulation-on* and *stimulation-off* periods in both control and targeted animals (**Supplementary Figure 3F**). We also observed that theta oscillations continued to remain disrupted during *stimulation-off* periods on the W-track. Thus, the disruption of theta during *stimulation-on* periods leads to disruptions of the fine-timescale temporal structure throughout the W-track experiences while leaving place fields unaffected.

These findings motivated a population-level approach to better understand the impact of theta disruption on representations of position on the W-track. We used a clusterless state-space decoding algorithm to estimate, for each 2 ms bin, the probability distribution of represented locations on the track (Denovellis et al., 2021).

Typical patterns of spatial representation were readily visible in control animals. There was pronounced rhythmicity reflecting theta sequences: each theta cycle typically begins with a representation close to the animal which then sweeps forward, resetting once the next theta cycle begins (**Figure 4A**) (Johnson and Redish, 2007, Foster and Wilson, 2007, Kay et al., 2020, Joshi et al., 2023). We quantified the prevalence of theta-timescale position sweeps by first calculating the distance (in 2 ms bins) between the peak of the posterior of the decoded representation and the animal’s actual position (ahead/behind distance). We then computed the power spectrum of this distance separately for outbound and inbound trials, given the distinct cognitive demands associated with each trial type. In the control animals, this power spectrum shows the expected pronounced peak around 8 Hz in both *stimulation-on* and *stimulation-off* periods (**Figure 4B**). We also quantified the changes in the probability of observing continuous decodes relative to stimulation times during *stimulation-on* periods. These probabilities showed the expected decrease in continuity associated with late phase theta, where the representation can jump back to a location at/or behind the actual location of the animal (**Figure 4C**). Similarly, the peri-stimulation patterns of change in entropy, which measures the concentration of the decoded distribution, showed the expected increase seen during late phase theta where spatial representations are less concentrated (**Figure 4D**).

**Figure 4.**
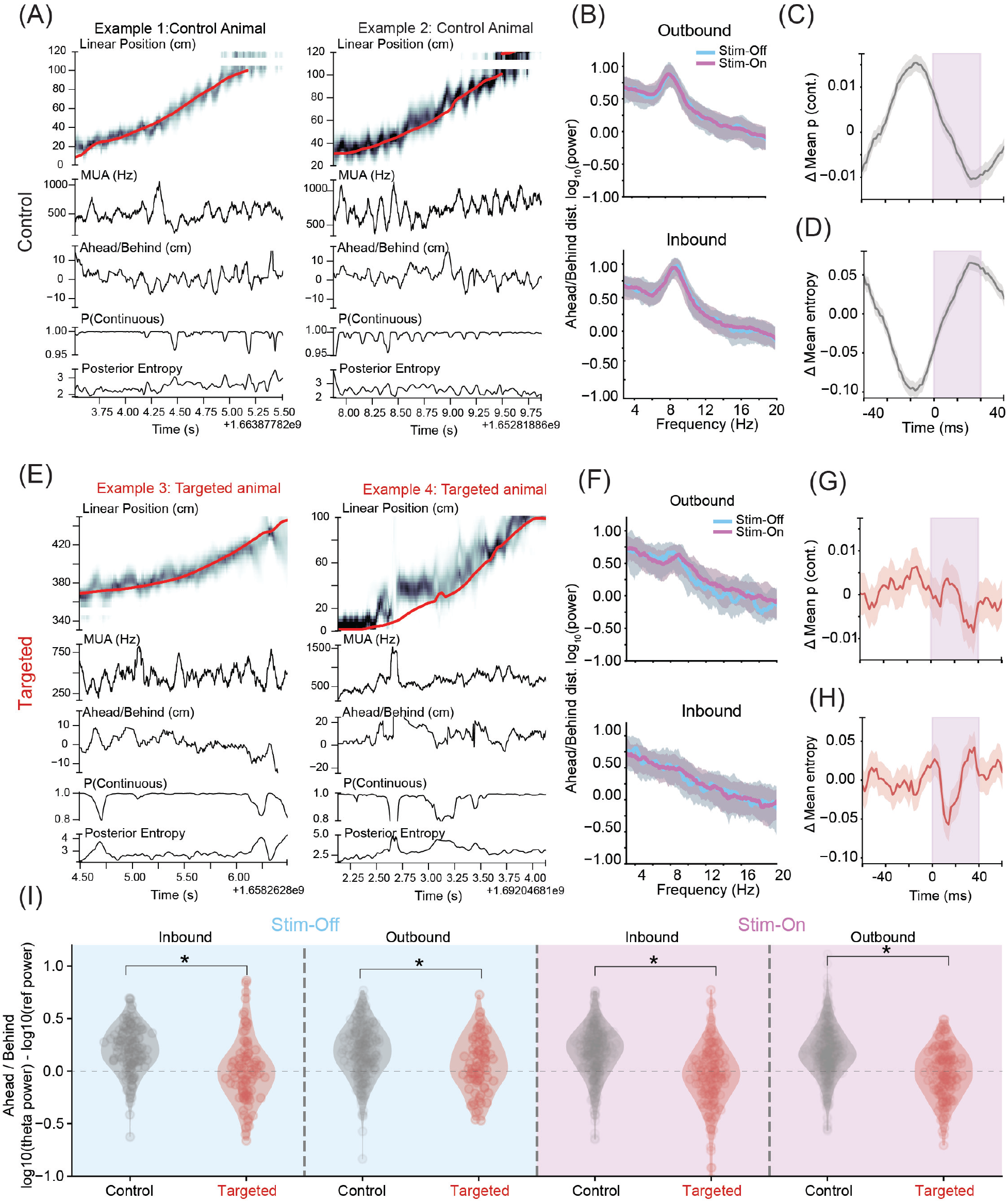
Theta disruption suppresses fast timescale sequential position representations. **A**, Examples of decoding position from population spiking on outbound trials in control animals. From top to bottom: actual linearized position (red) and estimated position posterior (greyscale); multiunit spike rate (spikes/s); ahead/behind distance of decoded position to animal’s actual position (cm); probability of a spatially continuous decoding state; and entropy of posterior over space. **B**, Power spectrum of the ahead/behind distance for *stimulation-on* and *stimulation-off* intervals across all trials in control animals, separated by trial type (top: outbound; bottom: inbound). Note the clear peak near 8 Hz reflecting the presence of robust theta sequences. **C**, Quantification of stimulus-triggered probability of the posterior to be continuous during *stimulation-on* trials for control animals. Note prominent theta-timescale structure reflecting more continuous representations in early phases of theta. **D**, Quantification of stimulus-triggered posterior entropy across all trials in control animals. Note increases in entropy toward later phases of theta corresponding to stimulation times. **E**, Examples of decoding position from population spiking on outbound trials in targeted animals; plot elements as in **A**. Note the disruption in theta-timescale structure. **F**, Power spectrum as in **B** for targeted animals. Note suppression of theta-timescale peak on outbound trials (top) and absence of obvious structure on inbound trials. **G**, Probability of continuous state as in **C** for targeted animals. Note altered distribution as compared to **C**. **H**, Entropy of posterior as in **D** for targeted animals. Note altered distribution as compared to **D**. **I**, Quantification of relative power in the power spectrum of the ahead/behind distance in the 8–12 Hz band relative to the mean of the 4–8 and 12–15 Hz bands. All comparisons were significant (all pooled data pairwise *t*-tests: *p <* 10*^−^*^4^; all hierarchical bootstrap tests: *p <* 0.05).

The theta-timescale rhythmic features of the population representation of space were disrupted in targeted animals, even as the decode remained on average close to the animal (reflecting the preservation of place field structure) (**Figure 4E**). Moreover, the rhythmic features were disrupted during both *stimulation-on* and *stimulation-off* periods. The *∼*8 Hz peak in the power spectrum of the ahead/behind distance was much less visible on outbound trials and was absent on inbound trials (**Figure 4F**). Further, the consistent modulation of both the probability of a continuous decode state and of the entropy of the posterior distribution around stimulation times was also much less prevalent in targeted animals (**Figure 4G, H**).

We focused on the power spectrum to quantify these differences across control and targeted animals, across outbound and inbound trials, and across *stimulation-on* and *stimulation-off* periods. Consistent with the pairwise analyses (**Supplementary Figure 3B-D**), the *∼*8 Hz peak in the power spectrum of the ahead/behind distance was significantly larger in control animals than in targeted animals across all conditions (**Figure 4I**; pooled comparison p’s < 10*^−^*^4^, hierarchical bootstrap p’s < 0.05). At the same time, maximal extent of locations represented ahead and/or behind the animal near the choice point, where choices must be made on outbound and inbound trials, did not differ between control and targeted animals (**Supplementary Figure 3G**). Thus, our findings indicate that the precise timing of non-local representations during theta was disrupted, but the spatial extent was not.

Finally, the differences in the power spectrum also motivated us to return to the linear track data and examine the impact of rhythmic frequency stimulation on decoded representations. To our surprise, the peak of the power spectrum of the ahead/behind distance appeared to shift systematically with the applied stimulation frequency (**Supplementary Figure 4**). This suggests a surprising level of preserved representational movement across frequencies ranging from 6 to 12 Hz. We also observed that in a subset of neurons, phase precession could be observed even when evaluated against the entrained *∼*10 Hz LFP (**Supplementary Figure 5**).

### Theta disruption spares replay during sharp wave ripples

A previous study that used an optogenetic intervention in the Entorhinal Cortex to disrupt hippocampal theta sequences reported that doing so also disrupted the sequential structure of replay events seen during subsequent offline periods (Liu et al., 2023). We therefore asked whether our manipulation could provide a dissociation between theta sequences and subsequent replay, or whether both were affected.

We first focused on replays seen during pauses in behavior on the track and found that despite the temporal disruption of spiking in the hippocampus during locomotion, awake SWRs and associated replays were not detectably affected. Replay events in both control and targeted animals showed the expected continuous progression of positions over time, along with increases in multiunit firing rates and power in the ripple band (**Figure 5A**). Moreover, we were able to identify clear replay events during the stimulation period within the first exposure to the track in both control and targeted animals (**Figure 5A** bottom row).

**Figure 5.**
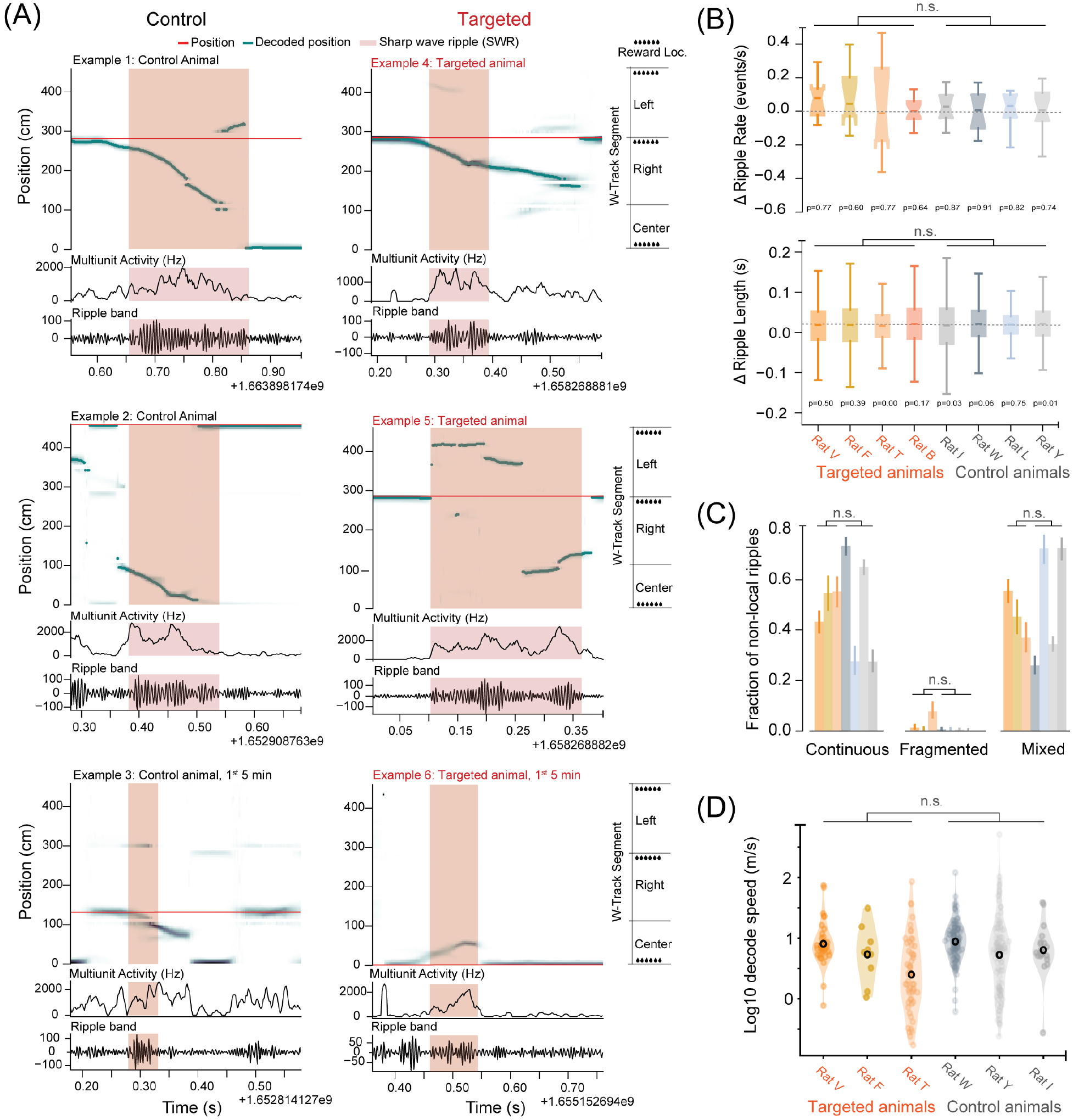
Theta disruption does not impact awake sharp-wave ripples (aSWRs) or replay. **A**, Examples of decoding position from population spiking in control (left) and targeted (right) animals during aSWR events. Bottom row shows events occurring during the first five minutes of experience on the track (a *stimulation-on* period). Each panel: Top: actual linearized position (red) and estimated position posterior (greyscale). Middle: multiunit spike rate (spikes/s). Bottom: bandpass-filtered ripple events. SWR events highlighted in light pink. **B**, SWR rates (top) and length (bottom) do not differ significantly between *stimulation-on* and *stimulation-off* intervals in targeted (shades of orange) and control (shades of grey) animals (paired *t*-test for each animal, *p* values reported in the figure; comparison of groups not significant, mixed effects linear model *p* = 0.929 for rates, *p* = 0.426 for lengths). **C**, Fractions of non-local SWRs that are spatially continuous, fragmented, or a mix of continuous and fragmented states, in targeted (orange) and control (grey) animals. Note overlapping distributions for each ripple content category. **D**, Average movement speed of the peak of the posterior distribution for continuous decodes (duration *>* 50 ms, probability of the event being continuous *>* 90)%. Distributions of speeds were very similar across control and targeted animals (mixed effects linear model, *p* = 0.253). Note, Rat B had a 32Ch linear probe and hence could not be used for SWR-associated content analysis. Rat L did not have enough continuous replay qualifying the inclusion criteria, but relaxing the inclusion criteria yielded similar results. We noted that there were some very high speed values. A visual inspection of those events suggested that they arose from cases where the posterior distribution was bimodal, and the peak of the distribution transitioned from one mode to the other over a single timestep, thus suggesting high speed movement even though each mode of the distribution was evolving much more slowly.

We then quantified SWR and replay properties based on three parameters that have been linked with hippocampal-dependent learning: SWR rate (Ego-Stengel and Wilson, 2010), duration (Fernández-Ruiz et al., 2019, Vancura et al., 2023), and content (Pfeiffer and Foster, 2013). There were no consistent changes in the SWR rate and length between control and targeted animals (**Figure 5B**). Next, we evaluated whether the content of the replay of neural activity may be altered in targeted animals. Here again, we found no evidence that theta disruption affected replay structure: sequential replays reflecting non-local spatial locations were observed during awake immobility with similar frequency across control and targeted animals (“continuous state,” **Figure 5C**). We then measured the speed of representation movement across >50 ms long continuous periods (**Figure 5D**). These speeds showed the expected concentration around 10 m/s (Davidson et al., 2009) and were not different across control and targeted animals.

Finally, we repeated a subset of these analyses during the rest sessions before and after W-track experience. Here again we were able to identify clear sequential replay events (Supplementary Figure 6A). We also found no differences in ripple rate or length across targeted and control animals (Supplementary Figure 6B). Similarly, we found that sequential replays were observed equally frequently across animals from both groups (Supplementary Figure 6C), and that there was also the expected increase in ripple rate from the first rest epoch, which preceeded the first run epoch of the day, and subsequent rest epochs (Supplementary Figure 6D). Thus, our manipulation specifically disrupted the temporal order of neural activity in single CA1 cells and their sequential population representations during locomotion without having a detectable effect on SWRs or replay.

## Discussion

We developed a targeted theta-phase-specific manipulation of hippocampal neural dynamics that selectively impacted sequential activity during locomotion while preserving immobility-related neural dynamics. We found that activating medial septal PV+ neurons at the ascending phase of theta during locomotion disrupted pairwise temporal properties and population-level spatial representations in the hippocampus, yet maintained the individual cell place code. This manipulation, applied during the first and final third of each run epoch, led to a profound impairment in learning the more cognitively demanding component of a hippocampal-dependent task while sparing learning on trials where a less cognitively demanding choice was required. Population neural activity patterns during immobility remained intact, indicating that locomotion-associated theta oscillations and immobility-associated replay mechanisms are functionally separable. Notably, despite this preservation of replay, learning deficits were as severe as those observed in a complete hippocampal lesion (Kim and Frank, 2009).

Our results extend findings from prior manipulations of hippocampal neural activity via medial septal manipulation (Wang et al., 2015, Fuhrmann et al., 2015, Robinson et al., 2016, Zutshi et al., 2018, Kloc et al., 2020, Mouchati et al., 2020, Quirk et al., 2021, Etter et al., 2023, Gemzik et al., 2021). While previous studies measured the impact of septal manipulation on theta, these studies did not systematically examine the fine timescale structure of population representations. Here we find clear evidence that driving the septal parvalbumin neurons at requested frequencies (rhythmic stimulation condition) drives hippocampal spatiotemporal sequences at that requested frequency. This finding is broadly consistent with the results of medial septum cooling, which slows both theta and the speed of progression of spatial representations in each theta cycle (Petersen and Buzsáki, 2020).

We also found a profound learning impairment for the outbound trials of our alternation task in an initially novel environment. This complements the demonstrations of impairment in performance of a learned task in previous studies (Zutshi et al., 2018, Quirk et al., 2021, Etter et al., 2023, Petersen and Buzsáki, 2020). In our work, we extend those studies to show that SWR rate, length and content are not impacted by the specific theta disruption paradigm we employed.

The importance of precise temporal organization of spiking during theta for learning is consistent with the hypothesis that this organization facilitates synaptic plasticity (Skaggs et al., 1996, Lisman and Jensen, 2013, Buzsáki, 2002). Our results provide strong support for this hypothesis: disruption of theta sequences was sufficient to profoundly impair learning on outbound trials while having no detectable effect on average place field properties or on learning on the interleaved inbound trials. Interestingly, recent studies from disease models also show disrupted synaptic plasticity accompanied with impaired theta sequence coordination and spared individual place fields (Viana da Silva et al., 2024, Donahue et al., 2025) suggesting that theta temporal structure is particularly vulnerable to plasticity disruption across both experimental and pathological contexts.

The dissociation between outbound and inbound learning mirrors patterns seen with awake SWR disruption (Jadhav et al., 2012) and indicates that disrupting either pattern is sufficient to lead to a learning deficit, although theta disruption appears to generate a more profound effect. The selectivity to outbound trials also offers clues as to the computations that may be disrupted. Outbound trials require that animals remember the outer arm they came from on the previous inbound trial such that they can use that information to choose to go to the opposite outer arm. This requires associations across locations, and correctly timed theta sequences may be critical for learning and retrieving these associations. Specifically, Hasselmo (2025) proposed that early phases of theta support information encoding, while later phases support retrieval (Hasselmo et al., 2002, Hasselmo, 2025, Mizuseki et al., 2009). Consistent with this hypothesis, hippocampal theta sequences typically represent locations close to the animal during early phases of theta and can represent hypotheticals, including possible futures, goals and alternative pasts, during later phases (Comrie et al., 2022, Foster and Wilson, 2007, Yu et al., 2026, Tang et al., 2026, Kay et al., 2020, Wikenheiser and Redish, 2015). The possibility that the timing of these representations is critical for normal brain function is consistent with our observation that representations of locations more distant from the animal were preserved during theta disruption, but the timing of these representations was no longer regular. Thus, the reliability and rhythmicity of septal input to the hippocampal formation are likely critical for providing temporal windows in which relevant behavior-related patterns can be organized into learning and action for outbound trials (Joshi et al., 2023, Joshi and Somogyi, 2020, Forli et al., 2025).

By contrast, a simple location-action association is sufficient for inbound trials, where animals could learn to execute a left or right turn when coming from the right or left outer arms, respectively. Thus, preserved place field activity in targeted animals could enable normal inbound learning. This specific impairment suggests a role whereby theta sequences are critical not only for goal representations (Yu et al., 2026, Tang et al., 2026) but additionally during cognitively relevant task phases (Joshi et al., 2023, Comrie et al., 2024).

Critically, the behavioral effects in targeted animals were seen even though stimulation was off during the middle third of each exposure to the W-track. Consistent with this behavioral result, sequential firing during locomotion (at both the pairwise and population level) was disrupted during *stimulation-on* periods and remained disrupted in *stimulation-off* periods, indicating that the 5-6 minutes of stimulation-off trials was not sufficient to allow the system to recover. This surprising result indicates that the disruption of theta sequences during the early experience in a novel environment is sufficient to have lasting effects, potentially by interfering with the rapid plasticity and place field formation induced during initial exposure. In this framework theta sequences would be particularly important for establishing task-relevant structure during the earliest phases of exploration. We also note that we can not rule out the possibility that a locomotion-specific incomplete physiological recovery from the preceding stimulation on the w-track may additionally contribute to the learning deficits. While we did not explicitly test the effects of pretraining or longer duration of stimulation-off periods, our results raise the possibility that pretraining the animal in the behavioral arena may allow for the development of task relevant representations, and thereby reduce or eliminate the behavioral impact of theta disruption.

Understanding precisely why this temporal organization is critical will require more distributed measurements. Notably, MS targets include multiple cortical and subcortical targets (Joshi, 2017), and our and previous manipulation may have disrupted precise spike timing throughout these regions. Key amongst these regions include the pre- and para-subiculum, retrosplenial area and the entorhinal cortex, which also receive dense PV projection in addition to the CA3 and DG (Joshi et al., 2017, Viney et al., 2018, Salib et al., 2019). The para-subiculum and entorhinal cortex (EC) also express sequential activity in the theta frequency range (Vollan et al., 2025). Disrupting the spatial code or spike-timing in these regions may contribute to the disruption of sequential activity we have observed. Additionally, monitoring regions beyond the temporal cortex would be informative given the broad coordination between hippocampal theta and other systems (Joshi et al., 2023, Eichenbaum, 2017, Buño and Velluti, 1977, Berg et al., 2006, Ledberg and Robbe, 2011).

Our results also demonstrate that theta sequences and replay sequences are dissociable. While our manipulation disrupted theta sequences on the track, we saw no effect on replay during periods of immobility on the track. Instead, we observed very similar patterns of activation of sequential representations during SWR events in targeted and control animals. This result contrasts with the conclusion of a previous paper (Liu et al., 2023) that showed the disruption of hippocampal theta sequences and subsequent replay during rest using an optogenetic manipulation of the EC. While our study and the previous study both employed disruptions limited to locomotion, one possible explanation for the difference is that there may have been sufficient recovery of sequential activity during stimulation-off periods to support the development of replay sequences, even though this activity remained disrupted as compared to controls. Previous results established that a single pass through a novel linear track is sufficient to enable replay (Foster and Wilson, 2006) and thus even a single pass where theta sequences had recovered could be sufficient to enable the expression of subsequent replay. In any case, our results show that replay alone is not sufficient to enable learning. Replay was intact while neural dynamics during locomotion were impaired, and this manipulation was accompanied by a critical impairment in learning the more cognitively challenging outbound component of the W-track.

Our findings, and previous dissociations between precise timescale and place field properties (Petersen and Buzsáki, 2020, Liu et al., 2023, Wang et al., 2016) further suggest that different circuits with different time constants are responsible for processing spatial and temporal information in the hippocampal circuit. Spatial/contextual information may arrive from regions with slower timescales (such as the cortex), making them less susceptible to sub-second brief disruptions, while precisely timed inputs from the medial septum coordinate the tightly controlled timing offsets between hippocampal neurons. We hypothesize that learning requires the intersection of these two streams of information in the hippocampal network, and is impaired by the disorganization of the precise temporal templates in which internal plans can be matched to external inputs.

Gaining a deeper understanding of the roles of these inputs will depend on manipulations that target specific patterns of organization of neural activity. This work will build on a wide array of previous studies that developed approaches to disrupt or alter pattern-specific firing (e.g., in medial septum (Wang et al., 2015, Fuhrmann et al., 2015, Robinson et al., 2016, Zutshi et al., 2018, Kloc et al., 2020, Mouchati et al., 2020, Quirk et al., 2021, Etter et al., 2023, Gemzik et al., 2021), entorhinal cortex (Liu et al., 2023, Lepperød et al., 2021), nucleus reuniens (Ito et al., 2015), supramammillary nucleus (Farrell et al., 2021), and pedunculopontine nucleus (Kaur et al., 2025)). We suggest that greater incorporation of open and closed-loop feedback paradigms will enable a better dissection of the roles of specific patterns of input and information flow in these circuits.

In summary, our results establish a dissociation between locomotion-related theta sequences and offline replay during learning, and indicate that disrupting theta seqeunces during locomotion is sufficient to drive a profound and specific learning deficit. This points toward a critical role of theta-sequences in building and accessing representations that enable choices that depend on integrating past and potential future locations.

## Acknowledgments

This work was supported by grants from the Simons Foundation 1189761, 1283731 (Joshi), 542981 (Frank), Life Sciences Research Foundation Fellowship to Joshi, NSF GRFP 1650113 (Comrie), NIH F31MH124366 (Comrie), UCSF Jonas Cohler Discovery Fellowship (Comrie), National Institute of Mental Health grant F30MH126483 (Guidera), National Institutes of Health grant T32GM007618 (Guidera), UCSF Discovery Fellowship (Guidera), and Phi Beta Kappa Graduate Award (Guidera), and funding from the Howard Hughes Medical Institute (Frank). We would like to thank Megan Carey and lab members for fruitful discussions and hosting Joshi during the analysis tenure of the project. We would also like to thank Anna Gillespie, Eleonore Duvelle, Gyorgy Buzsaki, and David Redish for comments on a previous version of the manuscript; Daniel G. for assisting in multi-electrode implant building; Erik H. for assisting in DLC model labeling; Trevor N. for assisting in data collection.

## Author Contributions Matrix

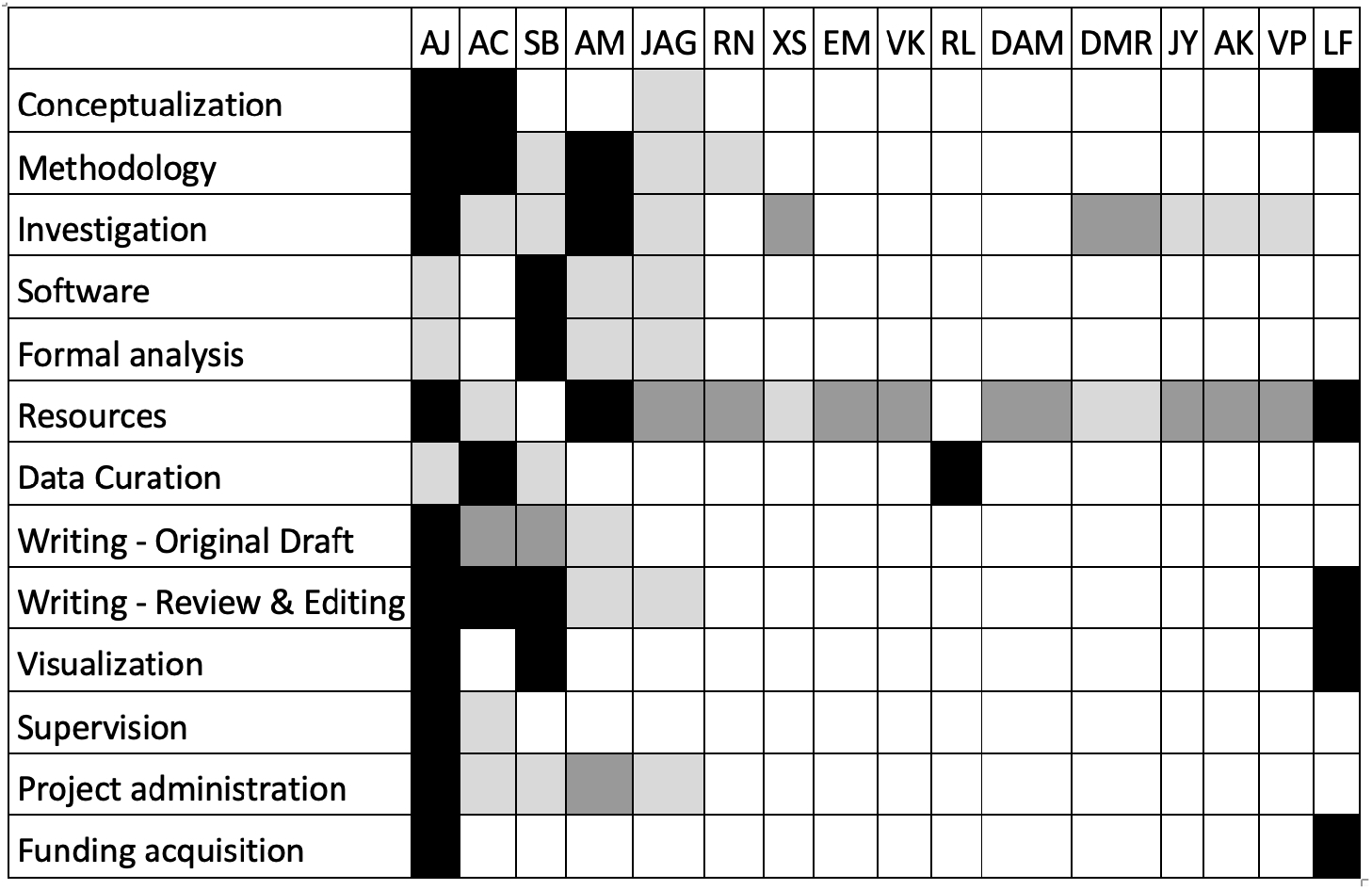

## Supplementary Information

**Figure S1.**
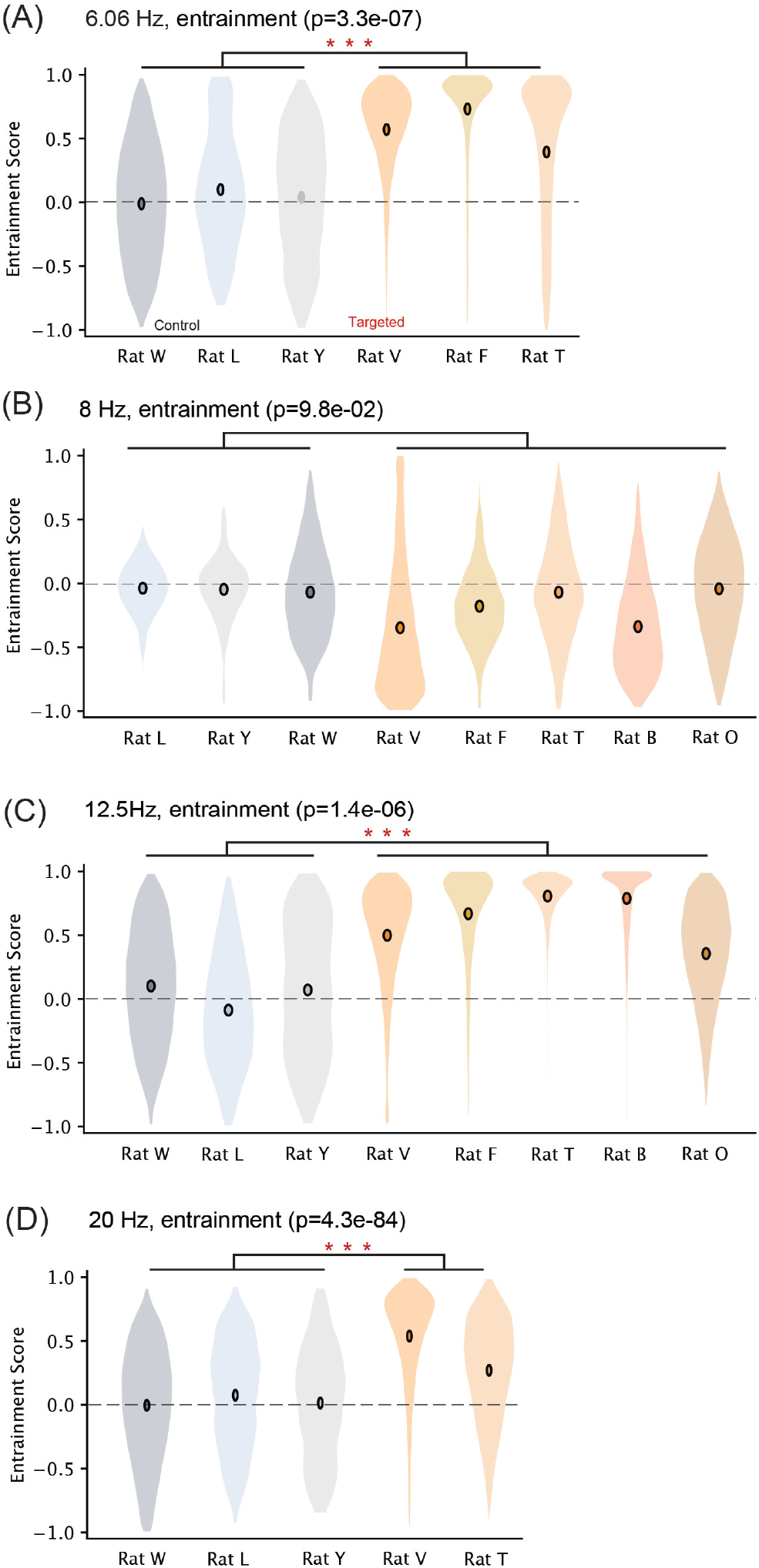
Entrainment of theta at different stimulation frequencies. **A**, Entrainment scores distributions across sessions for control (*n* = 3, grey) and targeted (*n* = 3, grey) animals at 6 Hz entrainment frequency. Targeted animals show significant entrainment compared to controls (mixed effects linear model (animal, targeted state), effect of targeting; *p* = 3.3 *×* 10*^−^*^7^, control *n* = 17492 interval pairs, targeted *n* = 21040 interval pairs). **B**, Entrainment scores distributions across sessions for control (*n* = 3, grey) and targeted (*n* = 5, orange) animals at 8 Hz entrainment frequency (control *n* = 8012 interval pairs, targeted *n* = 34184 interval pairs). **C**, Entrainment scores distributions across sessions for targeted (*n* = 5, orange) and control (*n* = 3, grey) animals at 12.5 Hz entrainment frequency. Targeted animals show significant entrainment compared to controls (mixed effects linear model (animal, targeted state), effect of targeting; *p* = 1.4 *×* 10*^−^*^6^, control *n* = 20152 interval pairs, targeted *n* = 30655 interval pairs). **D**, Entrainment scores distributions across sessions for control (*n* = 3, grey) and targeted (*n* = 2, orange) animals at 20 Hz entrainment frequency. Targeted animals show significant entrainment compared to controls (mixed effects linear model (animal, targeted state), effect of targeting; *p* = 4.3 *×* 10*^−^*^84^, control *n* = 10315 interval pairs, targeted *n* = 12863 interval pairs).

**Figure S2.**
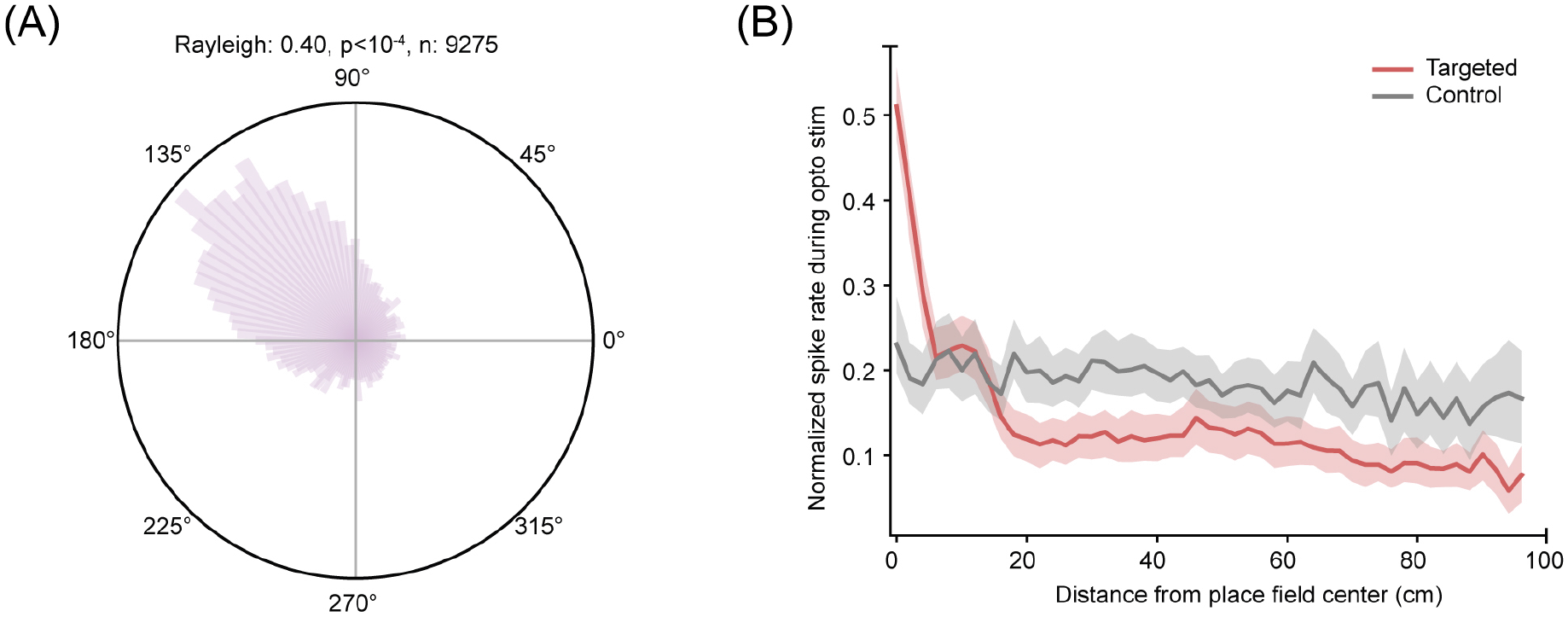
Accuracy and effects of theta phase–specific stimulation. **A**, Stimulus-triggered phase histogram for all linear track targeted animals (mean vector length = 0.4; *p <* 0.005; target phase = 90*^◦^*; pulse length *>* 20 ms; number of pulses = 9275). **B**, Normalized spike rate during optical stimulation in targeted (red, *n* = 307 neurons) and control (grey, *n* = 295 neurons) animals during theta phase–specific stimulation (theta phase = 90*^◦^*). For each neuron, spikes were collected within each stimulus event and binned by distance of the animal from the neuron’s place field peak defined from whole-epoch data. Shaded regions are the 95% CI with neuron resampling (**see Methods**). Note that extra spikes from stimulation are strongly localized to the place field center in targeted animals.

**Figure S3.**
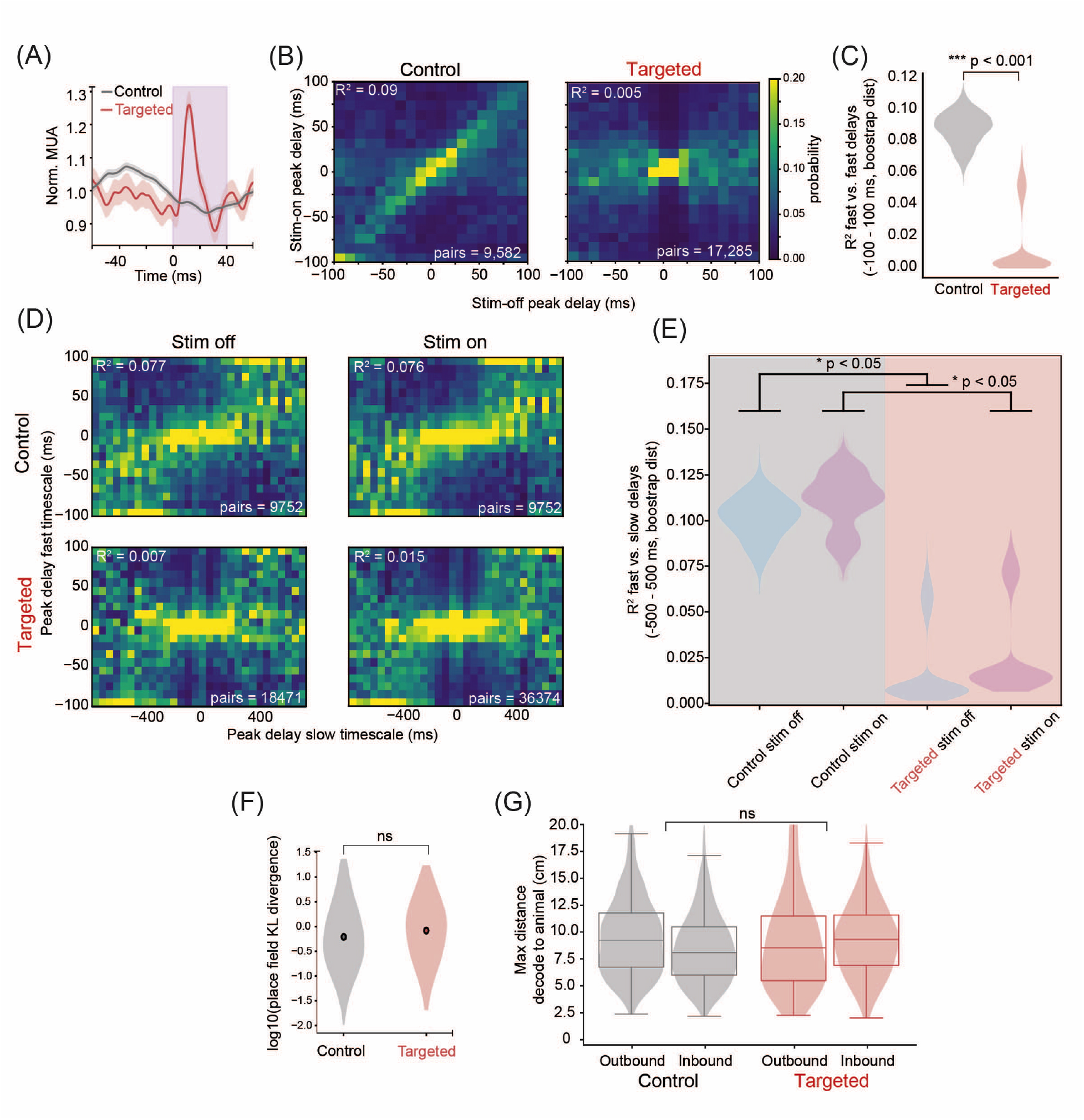
Disruption of short-timescale sequential spiking with preserved place fields in targeted animals. **A**, Quantification of normalized multi-unit activity triggered by onset of stimulation pulses shows an increase in spiking *∼*10 ms after stimulation onset on the W-track (as in the linear track) in targeted animals. As expected, control animals show a decrease in spiking (stimulation is phase-specific to the ascending phase of theta). Shaded areas are the 95% CI of the mean normalized firing rates in each condition. **B**, Pairwise temporal offsets between cross-correlograms of putative pyramidal neurons computed in *stimulation-on* versus *stimulation-off* intervals in control (*n* = 9582 pairs, left) and targeted animals (*n* = 17285 pairs, right). Note, a strong diagonal indicates that the cross-correlation structure is maintained in control animals (as expected). This structure is much less apparent in transfected animals during both *stimulation-on* and *stimulation-off* periods. **C**, Hierarchical bootstrap comparison of *R*^2^ values of linear fit, *p* = 8.6 *×* 10*^−^*^4^. **D**, Density plots of short- vs. long-timescale peaks for *stimulation-on* and *stimulation-off* periods for all cell pairs pooled across animals and run epochs. Top row: control animals; bottom row: targeted animals. The larger number of pairs for *stimulation-on* periods in targeted animals reflects a longer experimental period and stimulation driving larger numbers of short-timescale coincident firing events in cell pairs where too few events were detected for inclusion in the *stimulation-off* periods. **E**, Hierarchical bootstrap comparisons of *R*^2^ values across conditions demonstrating significant differences during both *stimulation-on* and *stimulation-off* periods. **F**, Place field stability for all units across control and targeted animals, measured as the KL divergence of their spatial firing selectivity between *stimulation-on* and *stimulation-off* intervals (*t*-test of distributions of KL divergence between targeted and control animals, *p* = 0.5). **G**, Quantification of the spatial extent of non-local representations at the choice point (maximum distance of the posterior *±*65 ms around the stimulus) during inbound (IN) and outbound (OUT) task phases on the W-track for targeted (red) and control (grey) animals. Note that both targeted and control animals have a similar extent of non-local spatial representations.

**Figure S4.**
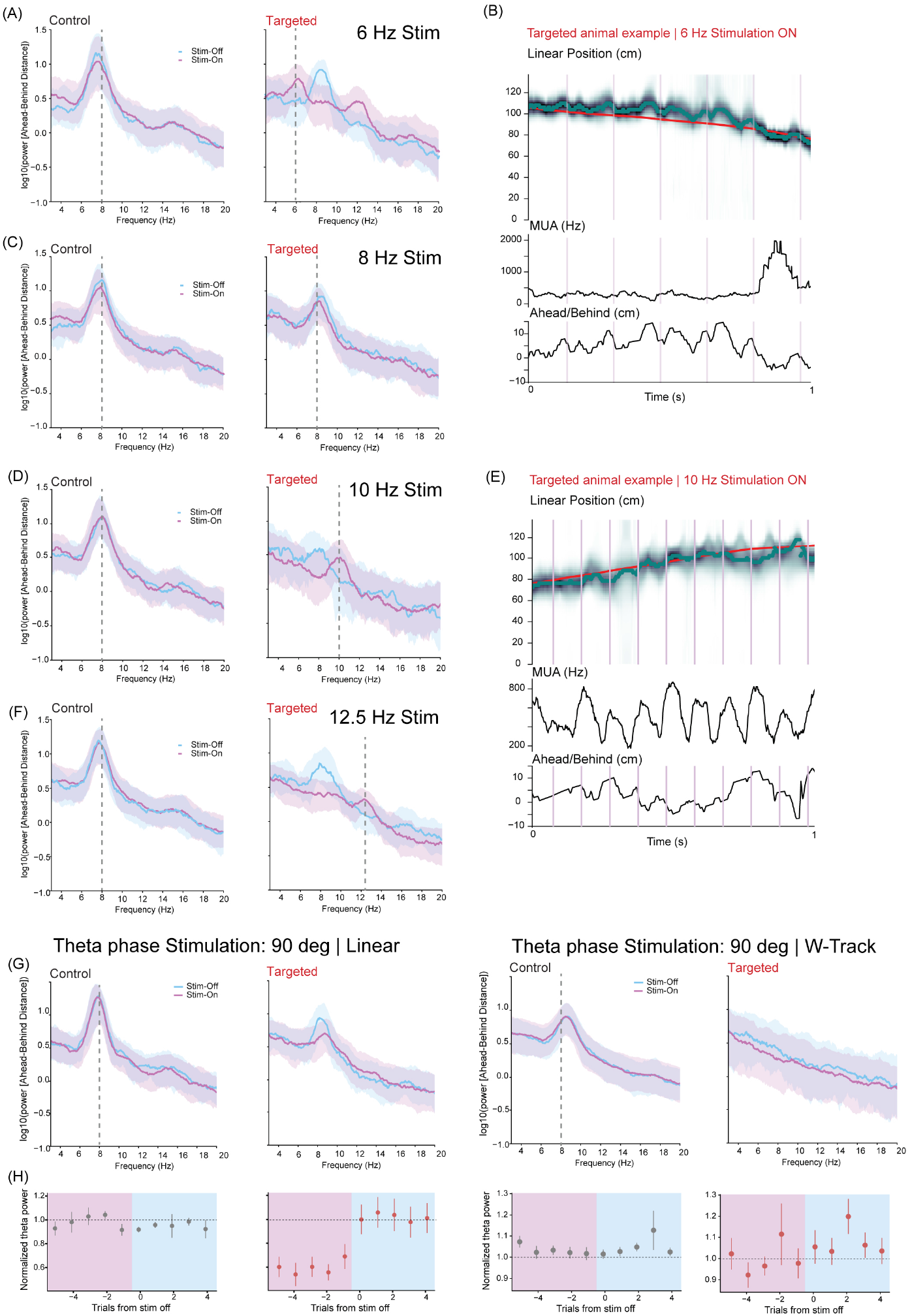
Effects of rhythmic vs. phase-specific stimulation on decoded representations. **A**, Power spectrum of the decode-to-animal distance trace for control (left) and targeted (right) animals in *stimulation-on* (purple) versus *stimulation-off* (blue) trials during rhythmic stimulation condition on the linear track. Note that the posterior rhythmically represents current and non-local positions at the targeted frequency of 6 Hz. Control animals continue to represent positions at 8 Hz despite the stimulation. This rules out that light flickering has an impact on the hippocampal representation. **B**, Examples of decoding position from population spiking during 6 Hz rhythmic stimulation on linear track. From top to bottom: actual linearized position (red) and estimated position posterior (greyscale); multiunit spike rate (spikes/s); ahead/behind distance of decoded position to animal’s actual position (cm). **C**, Power spectrum of the decode-to-animal distance trace for control (left) and targeted (right) animals in *stimulation-on* (purple) versus *stimulation-off* (blue) trials during rhythmic stimulation condition on the linear track. Note that the posterior rhythmically represents current and non-local positions at the targeted frequency of 8 Hz. **D**, Power spectrum of the decode-to-animal distance trace for control (left) and targeted (right) animals in *stimulation-on* (purple) versus *stimulation-off* (blue) trials during rhythmic stimulation condition on the linear track. Note that the posterior rhythmically represents current and non-local positions at the targeted frequency of 10 Hz. **E**, Examples of decoding position from population spiking during 10 Hz rhythmic stimulation on linear track. From top to bottom: actual linearized position (red) and estimated position posterior (greyscale); multiunit spike rate (spikes/s); ahead/behind distance of decoded position to animal’s actual position (cm). Note, that while our results indicate some movement towards the stimulated frequency, we cannot completely rule out the contribution of high spiking to the observed results. **F**, Power spectrum of the decode-to-animal distance trace for control (left) and targeted (right) animals in *stimulation-on* (purple) versus *stimulation-off* (blue) trials during rhythmic stimulation condition on the linear track. Note that the posterior rhythmically represents current and non-local positions at the targeted frequency of 12.5 Hz. **G**, *Left*, Power spectrum of the decode-to-animal distance trace for control (left) and targeted (right) animals in *stimulation-on* (purple) versus *stimulation-off* (blue) trials during theta-phase–specific stimulation targeted at the ascending phase of theta (90*^◦^*) on the linear track. Note that the posterior rhythmicity is markedly reduced in targeted animals compared to control animals. *Right*, Power spectrum of the decode-to-animal distance trace for control (left) and targeted (right) animals in *stimulation-on* (purple) versus *stimulation-off* (blue) trials during theta-phase–specific stimulation targeted at the ascending phase of theta (90*^◦^*) on the W-track. Note that the posterior rhythmicity is markedly reduced compared to control animals. We also note that the posterior rhythmicity does not recover during control periods on the W-track. **H**, Normalized theta power during trials after 90-degree theta phase stimulation was turned off on linear-track (left) and W-track (right) in control (gray) and targeted (red) animals. Note, almost complete recovery of theta power on the first trial after stimulation off on linear track and sustained reduction in theta power on the W-track.

**Figure S5.**
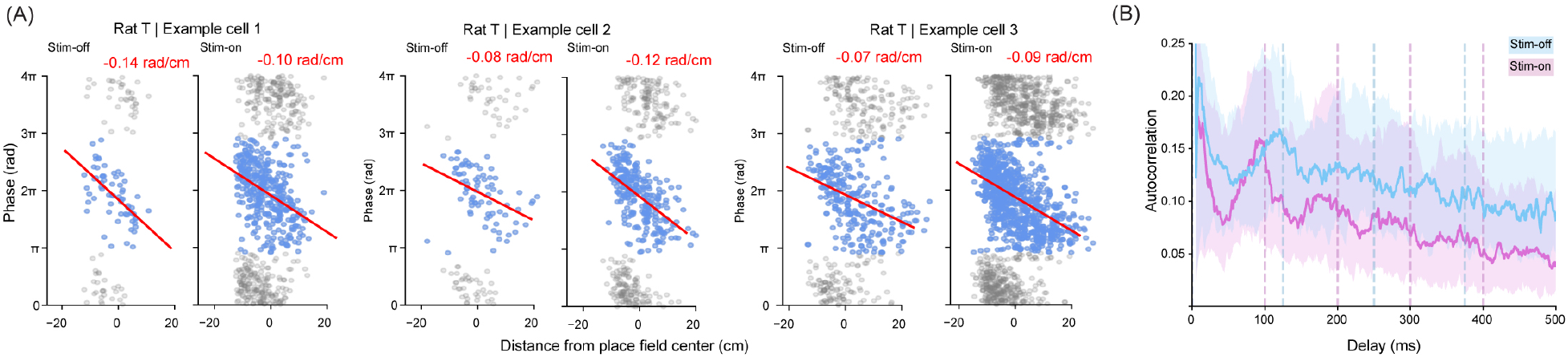
Theta phase precession in entrained LFP. **A**, Three examples of phase precession measured with reference to the stimulation off (endogenous theta) and stimulation on (entrained 10 Hz theta) showing sustained phase precession at the entrained theta frequency for individual neurons. **B**, At a population level, the autocorrelograms follow the entrained 10 Hz LFP in the stimulation-on condition compared to the stimulation-off condition (n=41 neurons, solid lines are medians, shaded areas 25/75 percentiles).

**Figure S6.**
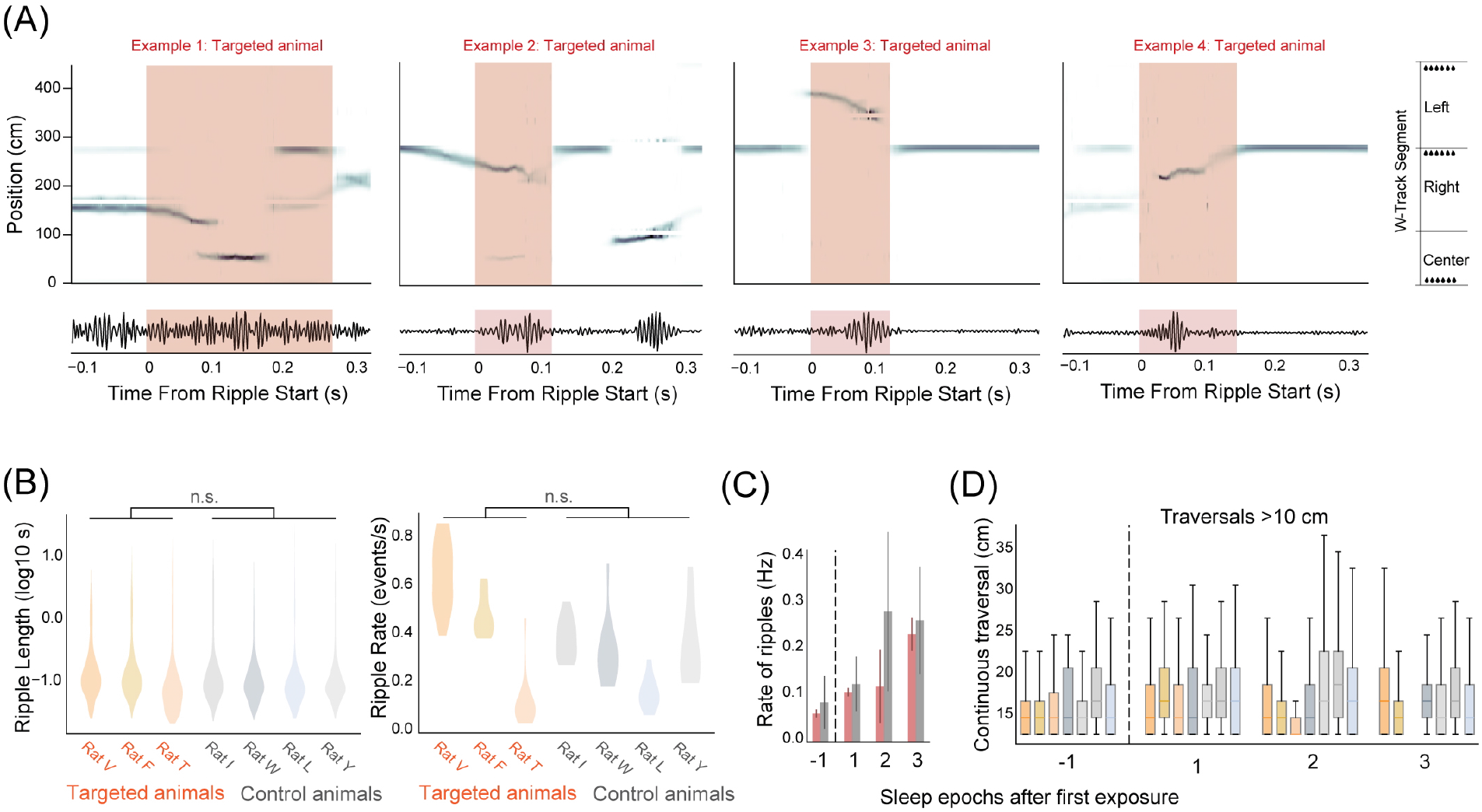
Theta disruption does not impact rest sharp-wave ripples (rSWRs) or replay. **A**, Examples of decoding position from population spiking in control (left) and targeted (right) animals during rSWR events. Top: estimated position posterior (greyscale). Bottom: bandpass-filtered ripple events. SWR events highlighted in light pink. **B**, rSWR durations (left) and rate (right) do not differ significantly between control and targeted animals (durations: p=0.94; rate: p=0.45; mixed effect model). **C**, rSWR rate increased in both control and targeted animals after exposure to a novel environment. **D**, Distribution of continuous traversals (>10 cms) for rest box epochs before and after the first exposure to the W-track.

**Figure S7.**
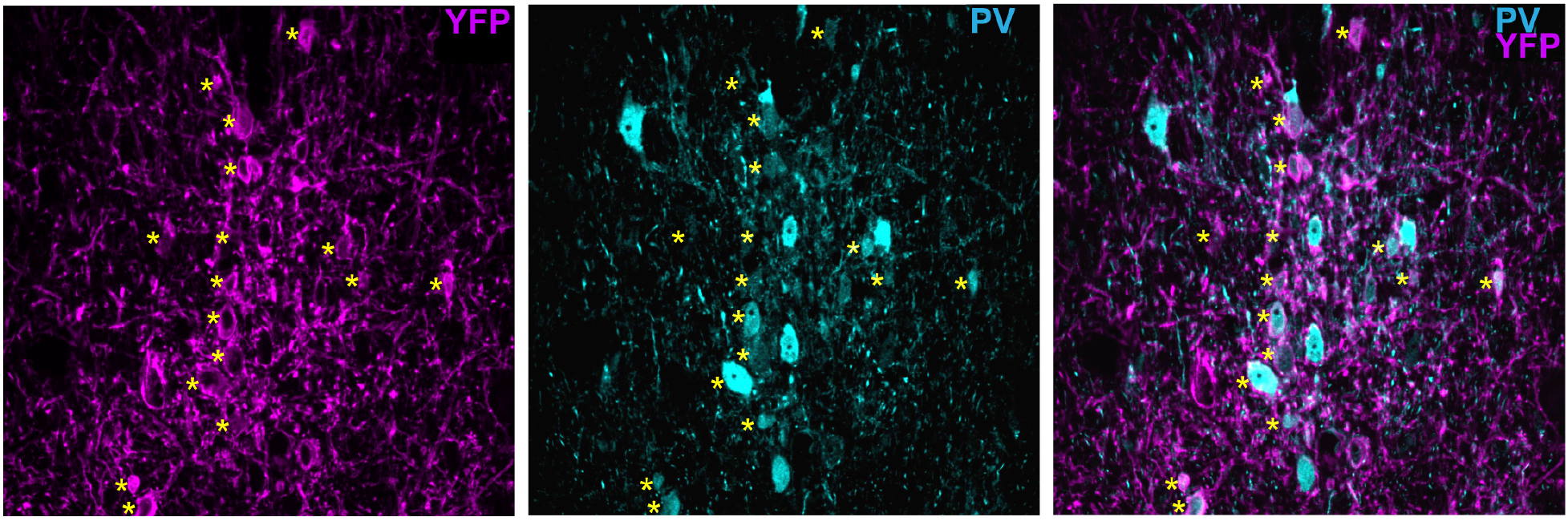
Overlap between eYFP expressing neurons and Parvalbumin in the medial septum. Injection site in the medial septum counter stained with anti-PV antibody (left), anti-eYFP antibody (middle), and merged images (right). Asterisks (yellow) mark eYFP neurons with eYFP expression in the cytoplasm.

**Figure S8.**
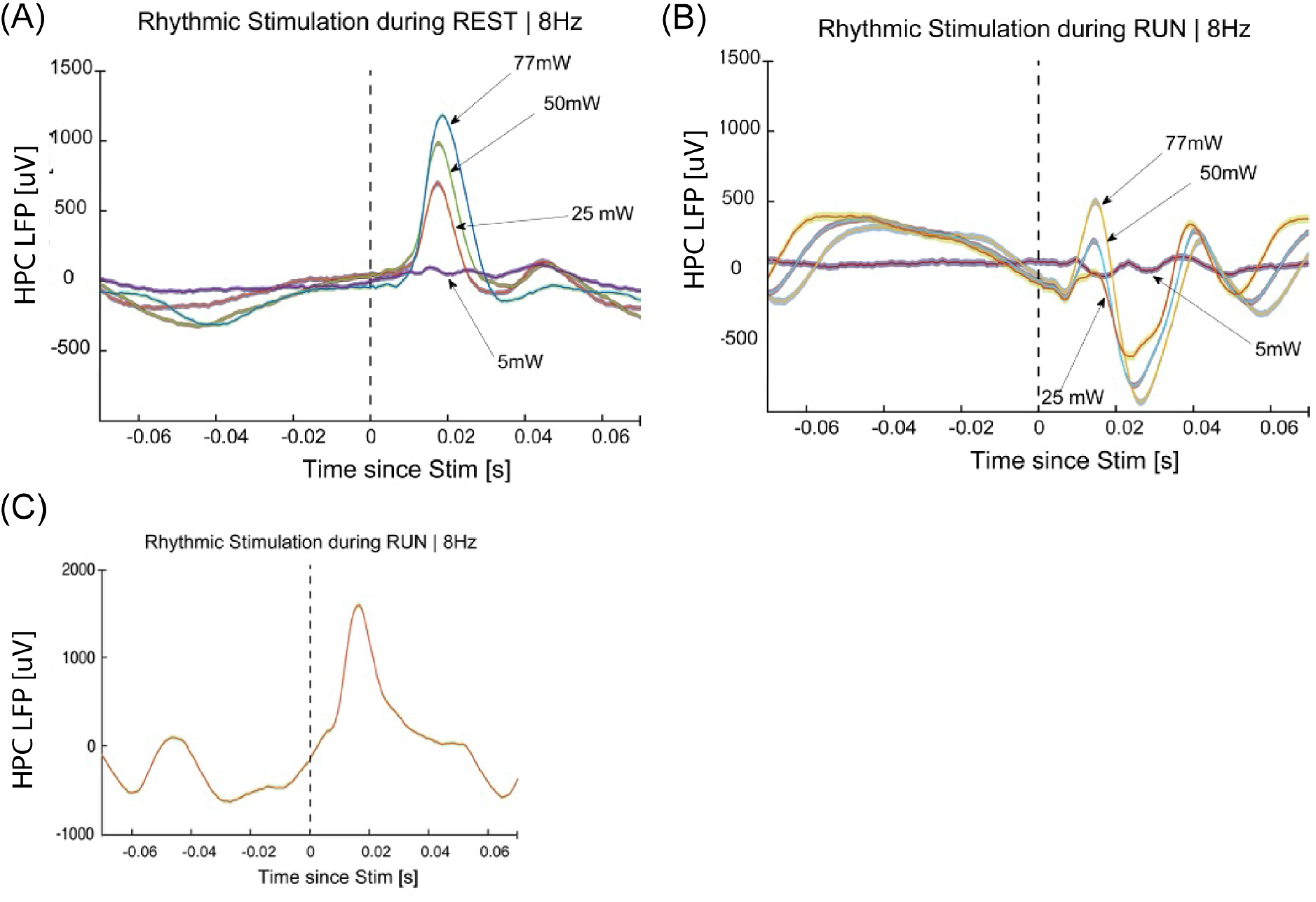
Laser-power dependence of LFP entrainment. **A**, Stimulus-triggered LFP in one targeted animal shows a laser power–dependent response during REST periods at 5, 25, 50, and 77 mW laser powers and 1 ms pulse width. **B**, Stimulus-triggered LFP in one targeted animal shows a laser power–dependent response during RUN periods at 5, 25, 50, and 77 mW laser powers and 1 ms pulse width. **C**, Stimulus-triggered LFP in one targeted animal at 77 mW laser power and 40 ms pulse width shows complete entrainment during RUN periods.

## Materials and Methods

### Experimental model and animal

Neural activity (cellular firing and local field potential) was recorded from the CA1 region of the dorsal hippocampus while simultaneously performing optogenetic stimulation of parvalbumin (PV) neurons in the medial septum in ten PV-Cre Long Evans male rats (6–10 months old, 500–650 g) (Yu et al., 2018). All experimental procedures were in accordance with the University of California San Francisco Institutional Animal Care and Use Committee and US National Institutes of Health guidelines.

### Surgery, virus injection, array insertion, and electrophysiological recording

Prior to surgical procedures, animals were maintained on full food (ad libitum) for one week. Surgery involved injection of a Cre-dependent virus (AAV5-Ef1a-DIO-hChR2(H134R)-eYFP, UNC Vector Core) into the medial septum (ML –0.15 mm, AP +0.904 mm, DV –6.94 mm) and implantation of a multi-electrode recording array in hippocampal area CA1 (AP –3.8 mm, ML +2.8 mm, DV set by tetrode adjustment). Virus (450 nL, 4 *×* 10^12^ viral particles/mL) was injected at 0.05 µL min*^−^*^1^ using a Hamilton Nanofil 10 µL syringe with a beveled 33G needle, mounted on a stereotaxic syringe pump (KD Scientific). After each injection, the needle was left in place for 5–10 minutes before retraction to ensure diffusion. Next, we inserted a tapered optic fiber (Optogenix, Lambda-B, NA = 0.39; core/cladding diameters = 200 µm/225 µm; emitting length = 2 mm; implant length = 6 mm; connector: 2.5 mm ceramic ferrule) following the same path as the injection needle. The optic fiber was inserted 200 µm deeper than the injection location to maximize lateral spread of the laser beam in the injection zone. After virus injection, rats were implanted either with 32-channel silicon probes (n = 3; NeuroNexus, 1 shank, 32 electrode sites, shank length = 6 mm, spacing = 50 µm, electrode surface area 177 *µ*m^2^) or with a 128-channel tetrode drive (v4.6). Animals were allowed 2–3 weeks for viral expression before experiments began. For the animals implanted with 32-channel silicon probes (*n* = 3), we tested the laser power and pulse durations required to entrain theta oscillations. All three animals were targeted as expected; however, for one animal, the maximum laser power tested was 50 mW, which was not sufficient to induce theta oscillatory activity during the run (only during rest periods). Hence, that animal was not included in the rhythmic stimulation analysis. For three animals, we assessed the fraction of eYFP expressing neurons that were parvalbumin expressing by co-staining medial septal injection sites with PV (anti-PV antibody; Sigma; P3088) and eYFP (anti YFP antibody; abcam; ab290). We found a high degree of overlap between eYFP positive cells and PV expressing neurons (greater than 75 percent (**Supplementary Figure 7**)) as expected from prior studies using the same rat strain (Yu et al., 2018, Lepperød et al., 2021). Each neuron was tested for parvalbumin expression in the cytoplasm and not in the nucleolus across multiple sections of imaging if the assessment could not be made in one imaging plane.

### Behavioral arenas

Rats were pre-trained on a linear track (125 cm long) to alternate between reward wells on either end. Prior to training, animals were food restricted to 85% of their starting body weight. Nose pokes at the wells were detected by an infrared beam, and upon a successful visit 200 µL of sweetened milk was delivered. Pre-training included two 30 min sessions interleaved with an hour of rest in the rest box.

#### Linear track

Animals implanted with the recording drives/probes were made to run on the linear track for sweetened milk reward while the electrophysiological signals were recorded using SpikeGadgets hardware and software (v.1.8.0). The animals implanted with the 128 Ch tetrode drives had to undergo a series of tetrode adjusting steps over 10–14 days to lower the tetrodes to the hippocampal cell layer before we started behavioral protocols. The recording setup involved multiple independently moving pulley systems running on carbon fiber tracks hanging from the ceiling, which supported the cables running from the animal’s head allowing the animals to move freely on the 2-D linear track plane. The connection from the drive was passed on through a rotating commutator before connecting to the computer allowing the animals to freely rotate their heads without putting rotational strain on the cables. The medial septal parvalbumin neurons were stimulated using an optic fiber from the laser connected to the optic fiber ferrule on the head of the animal. The animals underwent a stimulation with spatial constraint where the laser was ON only when the animals were away from the reward wells, thereby stimulating primarily during running times. The stimulation protocol (described below) involved pulses of varying width (pulse width values), intervals (interval values) and numbers (number of pulses).

#### W-Track

After linear track running sessions, animals ran on a W-shaped track to test spatial learning and memory-guided decision making. The track involved 3 arms with the sidearms connected to the base of the center arm forming a W-shape. The rule for this task was that the animals were rewarded when they alternate between the two side arms of the track while visiting the center arm in between each alternation. 0.2 mL sweetened milk was administered at the end of each arm for every correct visit. Electrophysiological data from dCA1 was recorded while the animals underwent a closed loop stimulation of the medial septal parvalbumin neurons. The closed loop stimulation involved both spatial constraint where the laser was ON only when the animals were away from the reward wells and theta phase constraint where the stimulation was tuned to the ascending phase of theta (90-degrees).

### Optogenetic manipulation setup

Optogenetic stimulation was achieved through an Omicron LuxX Plus laser (488 nm, 100 mW maximum output) connected to implanted optic fiber cannulae. The cannulae were purchased from Optogenix (Lambda-B, numerical aperture: 0.39, core/cladding diameter: 200 µm / 225 µm, emitting length: 2 mm, implant length: 6 mm, ceramic ferrule connector: 2.5 mm). Laser power was initially titrated, beginning at 5 mW and increasing gradually until reaching 77 mW, which was used for all subsequent experiments.

In the probe-only animals (*n* = 3), we tested the ability to entrain hippocampal LFP during the interleaving rest box periods, by varying laser power (from 5 mW to 77 mW at source) and pulse width (1–60 ms). We found that with a pulse width of 1–5 ms at laser powers greater than 25 mW, we could completely entrain hippocampal LFP in the rest box (when the animals are predominantly immobile), while higher laser powers (77 mW) and longer pulse widths (*>* 20 ms) were required for entrainment on the linear track (when the animals are engaged in the movement) (**Supplementary Figure 8**). Thus, for the rhythmic and phase-specific manipulation, we used 77 mW laser power at source and pulse widths greater than 20 ms. Note that while this is a high overall light power, the use of a tapered fiber substantially lowers the power density in the tissue as compared to a blunt fiber.

### Optogenetic stimulation protocols

Optogenetic stimulation protocols were delivered using a custom adaptation of the FSGui module of SpikeGadgets Trodes software. Publicly available code can be found at https://bitbucket.org/mkarlsso/trodes/branch/fsgui_theta2. This module enables two relevant modes of stimulation: rhythmic stimulation at a certain frequency, and theta-phase specific stimulation at the requested phase of the recorded theta rhythm. To limit stimulation to behavioral periods associated with locomotion and the associated hippocampal theta rhythm, we restricted use of these stimulation modes to periods in which the animal occupied a user-defined spatial area of the maze. Spatial filters were set to only permit stimulation when animals were locomoting on the maze, and to prevent stimulation when animals occupied reward port locations.

**Open loop stimulation** was delivered using the following parameters: Spatial lockout period in samples (set to 7500); Pulses per pulse (set to one for data presented in this paper); Period (ms): 50, 80, 100, 125, 165, 250; Pulse length (ms): 1, 5, 27, 40 (we resorted to using *>* 20 ms as it provided us with the best entrainment response as compared to 1 and 5 ms).

**Closed loop theta phase-specific stimulation** parameters enabled specification of a target theta phase. Briefly, one channel of 30 kHz of LFP data recorded from corpus callosum was downsampled to 1500 Hz. A second-order Butterworth filter with a short group delay was applied to the LFP data in the 4–12 Hz frequency band (similar to (Siegle and Wilson, 2014)). Upward- and downward-going zero-crossings were detected in the filtered data, and were used to estimate the period of the prior cycle. The period of the prior cycle and zero crossing times were then used to estimate the sample at which the incoming data would reach the target theta phase. A lockout period was specified to prevent excess stimulus triggers. This enabled stimulation at the target phase (Supplementary Figure 3). The closed loop stimulation parameters used in these experiments were: Target theta phase (90-degrees); Spatial lockout period in samples (7500); Theta lockout period (2400 samples); Pulses per pulse train (1); Pulse length (ms): either 27 (*n* = 3 probe animals) or 40 (*n* = 5 animals).

### Behavior analysis

A state space algorithm (Smith et al., 2004) was used to estimate the probability of a correct choice and accompanying 90% confidence interval on inbound, outbound, or all trials. The “learning trial” was defined as the first trial on which the lower bound of the 90% confidence interval was above chance (50%) (Smith et al., 2004). For each animal, we compute related behavioral metrics: the learning trial, the slope of the regression of performance on evenly spaced numbers from 0 to 1 equal to the total number of trials, and the number of trials performed correctly. We also computed average learning curves across animals within each group, and for each trial type, by finding the mean fraction of trials performed correctly in consecutive spans of 20 trials. We show these averages and accompanying 95% confidence intervals.

### Data analysis

#### Codebase and statistical approach

We prepared this dataset in a commonly accessible and reusable format, meeting the Neurodata Without Borders (NWB) standard (Rübel et al., 2022). This data was then ingested into the spyglass framework (Lee et al., 2024), which allowed for centralized data management, ensuring all intermediary steps from the raw NWB file to various analysis files were logged. Complete code for data processing and figure generation is available at: https://github.com/LorenFrankLab/ms_stim_analysis.

Across all experiments, an epoch is defined as a single period of time in the task arena, and is typically *∼*20 min. At most, one setting of the optogenetic stimulation protocol was run per epoch. Within each epoch, the *∼*5–7 min “*stimulation-off*” period during the middle of the epoch served as a control for within-epoch comparison of the effect of the driving stimulus. For most analysis, we restrict the data to the trial intervals, defined as the time between leaving a reward port to entering the next. Reward port occupancy was determined as the tracked head position being within 20 cm of the end of the track. For analysis of features specific to animal movement, these intervals were further restricted to times where the animal head speed exceeded 10 cm s*^−^*^1^.

Bootstrap confidence intervals were performed by resampling the observed distribution of events 1000 times and calculating the summary statistics for each. For hierarchical bootstrapping, both the animals within treatment groups and the events within animals were resampled for each calculation. *t*-test statistics were run using the scipy implementation. Mixed linear regression models were run using the statsmodels python package.

#### LFP analysis

LFP data was generated from the original 30 kHz electrophysiology recordings by filtering the data with a 0–400 Hz low-pass filter and downsampling to 1 kHz. Results were generated by analyzing the data from a reference electrode above the CA1 layer. Artifacts in the data were defined as periods where the amplitude of the filtered data exceeded 2 mV and were excluded from analysis.

LFP power spectrums were calculated from intervals when animals were outside the reward ports and traveling with a speed of at least 10 cm s*^−^*^1^. Power density spectrums and related scores described below were calculated for each continuous interval meeting these criteria using the scipy implementation of Welch’s method. These were then weighted by duration of the interval and averaged to produce the results shown. [Full analysis: func: `Analysis/lfp_analysis/opto_spectrum_analysis()`]

##### Entrainment score

Entrainment scores were calculated for open-loop fixed-frequency stimulations based on the power of the LFP spectrum at the target frequency during stimulus on (*p*_test_) and stimulus off (*p*_control_) intervals. [Full analysis func: `Analysis/lfp_analysis/calculate_entrainment_score()`.]

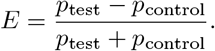

##### Suppression score

Suppression scores were calculated by first identifying the frequency with maximum power in the theta band (4 *< f <* 12 Hz) in the control power spectrum, then measuring power at that frequency in the control (*p*_peak_control_) and test (*p*_peak_test_) intervals. [Full analysis func: Analysis/lfp_analysis/calculate_suppression_score().]

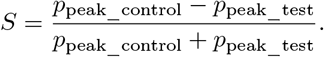

#### Spike Sorting

Spike sorting was performed in the `spyglass.spikesorting.v0` pipeline. Raw recording files were referenced and bandpass filtered from 600–6000 Hz. Artifacts in the recording were identified as periods where at least 75% of the filtered channel recordings exceeded 2000 µV with 500 ms padding on each side of the event. These were excluded from sorting. Spike sorting was run on each epoch through the spikeinterface package using mountainsort4 sorter with; signal whitening, adjacency_radius 100, clip_size 30, detect_threshold 3, and detect_interval 10. After sorting we computed quality metrics on the identified units using spikeinterface. We then excluded units with a nearest neighbors noise overlap greater than 0.03, a peak offset greater than 2, or an inter-spike interval violation greater than 0.0025.

#### Event-triggered firing rates for individual cells

For each sorted unit, spike events were binned based on time delay to the nearest stimulus onset. We then calculated the KL divergence between this distribution of spike delays and a null uniform distribution in the same interval. To determine if this represented a significant divergence from the null distribution, we performed circular shuffling of the stimulus times by offsetting each stimulus onset time by an independently chosen value from a range of *±* the stimulus period. The KL divergence between the shuffled delay distribution and the uniform distribution was calculated. This process was repeated 1000 times to form a null distribution of KL divergences for each unit. A unit was determined to be significantly modulated if the observed KL divergence was above the 99.5 percentile of the null KL distribution.

#### Event triggered MUA analysis

Multiunit firing rates were calculated by summing the total number of firing events from sorted units within 2 ms time windows, and then smoothing with a gaussian filter with standard deviation of 5 ms. The mean of this value at each timepoint across trials was estimated using bootstrap sampling techniques described above.

#### Place fields

Place fields were calculated by fitting a `SortedSpikesDetector` decoding model from the non_local_detector package (GitHub). Positions were first projected to the linear track skeleton to allow for 1D decoding. Decoding was performed using the packaged sorted spikes kernel density estimation algorithm with sampling rate of 500 Hz, position standard deviation of 6 cm, and discrete position binning of 2 cm. For comparisons of the place coding of cells between the optogenetic test and control intervals, we fit separate models using each of these segments as the encoding interval.

#### Unit place field coverage

For much of our analysis, we wanted to restrict to units which displayed spatially-localized distributions of firing activity. To do so, we calculated the highest posterior density of the place field distribution to determine the minimum spatial area required to capture 50% of this firing distribution. Thresholding an upper limit for this value allowed us to select cells with well-localized firing patterns.

#### Temporal offset of cross-correlation

Cross correlograms were calculated for intervals when the animal was continuously running outside of the reward port (speed *>* 10 cm s*^−^*^1^ for at least 500 ms). Histograms for all spike pairs were calculated over 1 ms time bins.

To calculate fast, theta-timescale delay times, the cross correlograms were then smoothed with a gaussian kernel with standard deviation *σ* =3 ms. Cell pairs were included for further analyses if they had at least 50 pairs of spike events co-occurring within a *±*100 ms window. To define the delay of peak cross-correlation times, we used the scipy function `find_peaks` with a distance equivalent to 80 ms and selected the delay peak with the largest magnitude.

For slow, positional timescale delay times, the cross correlograms were smoothed with a gaussian kernel with standard deviation *σ* =500 ms. Cell pairs were included if they had at least 100 pairs of spike events co-occurring within a *±*1000 ms window. Peaks were identified with a distance equivalent of 2 s, and the peak closest to zero taken as the delay time.

#### Place Field Similarity

Consistency of place field coding under the stimulus protocol was quantified by the KL divergence between the place distribution of each neuron’s firing during the control versus the test interval. Units were included in the KL-based place-field stability analyses only if they satisfied both of the following: **(i)** Spike-count criterion: at least 100 spikes in each interval type used for the comparison—*stimulation-off*, *stimulation-on*, and the *stimulus* sub-interval. **(ii)** Coverage criterion: the 50% highest posterior density (HPD) region of the unit’s spatial firing distribution was contained within 40 cm on the linear track or within 100 cm on the W-track.

#### Outbound repeat probability

Outbound repeat sequences are defined as the number of visits to the same outer arm without visiting the alternate arm within an epoch. Note that these sequences are typically interleaved with rewarded visits to the center arm. We show the distribution of sequences of length greater than *x* in Figure 3E. This distribution under random choice of outer arm at each trial is given as

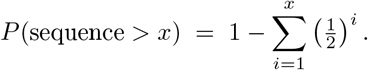

We sampled this random binomial choice process for the observed number of trials 1000 times for each animal group to get the 95% confidence interval of a random choice process in each case. Values for long repeat sequences below this interval indicate learning of the outbound task.

#### Clusterless Decoding

Clusterless waveform features were extracted from the same filtered recording used for spike sorting. Event times were identified using `spikeinterface.DetectPeakByChannel`, with an amplitude threshold of 100 µV and a local exclusion radius of 100 µm. Waveforms of each event were featurized by the peak amplitude within *±*500 µs of the event on each channel of the tetrode on which the event was detected. Clusterless position decoding was performed using the `ClusterlessDetector` algorithm from the `non_local_detector` package (GitHub). Positions were first projected to the linear track skeleton to enable 1D decoding. Decoding was then performed with the package’s clusterless kernel density estimation algorithm, using a sampling rate of 500 Hz, waveform feature standard deviation of 24 µV, position standard deviation of 6 cm, and discrete position binning of 2 cm.

As some animals had fewer detectable cells on later days of experiments, we restricted our subsequent decoding analysis to positions on the track with sufficient place cell coverage for reliable decoding. We defined this as track regions that fell in the 50% credible interval of at least two well-localized cells’ firing fields (95% of the units place field distribution localized to *<*50 cm of the track, see Unit place field coverage). From this we defined time intervals where the animal was physically located in these positions to use in decoding analysis below.

##### Ahead-Behind Distance

Orientation of the animals during movement was estimated using the arc tangent of the animal’s velocity. This value is reasonably well-defined during run intervals analyzed here. Distances between the position and decode along the track path are calculated using the `get_ahead_behind_distance` function in the `non_local_detector` package. Since our experimental paradigm suppresses theta oscillations themselves, we used a clusterless decoding approach (as in Joshi et al. (2023)) to obtain an unbiased estimate of rhythmicity for hippocampal spatial representations. Briefly, we estimated the peak of the posterior at every 2ms timestep and computed the distance between that value and the actual position of the animal (decode-to-animal distance). We confirmed that, as expected, in control animals, we could detect “theta sequences” as in prior studies without explicitly requiring theta oscillatory cycle windows and thus used the same method to detect hippocampal spatial representations in targeted animals.

##### Decoding Stimulus Response

Analysis was restricted to stimuli that occurred while the animal was running outside of the reward ports (speed *>* 10 cm s*^−^*^1^). Furthermore we excluded stimulus times with poor decoding (more than 10 ms of the *±*50 ms window surrounding a stimulus containing decodes *>*50 cm from the animal’s position).

##### Decode Coverage

Decode coverage is calculated as the area of the linearized track (in cm) spanned by the 95% credible interval of the position posterior from the decoding results.

##### Posterior entropy

Entropy of the posterior distribution over discrete position bins is defined as:

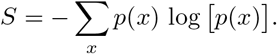

##### Ahead-behind distance

see above. Values are clipped to a range of 20 cm around the animal to prevent infrequent large events from skewing distribution.

##### Multi Unit Activity

Rate of spike events from all sorted neurons is calculated in 500 µs time bins and smoothed with a gaussian kernel of *σ* =1 ms.

##### Decoding Distance Power Spectrum

Intervals were identified where the animal was continuously running (*>*10 cm s*^−^*^1^) for at least 1 s outside of the reward port. Power spectrums were calculated for each of these intervals using the scipy implementation of Welch’s method with a 1 s window. The mean was taken as an average weighted by the length of each interval.

##### Decoding Distance Power Spectrum Theta Peak

For each spectrum calculated above, the average power in the theta range (6–10 Hz) was normalized by the average power in a reference frequency range (union of 2–6 Hz and 10–14 Hz). The theta peak of the spectrum was defined as the log of this ratio.

##### Decoding Ahead/Behind Maximum Distance

Intervals were identified where the animal was within a 20 cm radius of the choice point center. For each stimulus within these intervals we identified the maximum decode to animal distance with *±*65 ms of stimulus onset.

#### Sharp Wave Ripples

Sharp wave ripples were identified using the `spyglass.ripple.v1` pipeline. LFP data was first bandpass filtered in the range 150–250 Hz. Ripple times were then identified using the z-score threshold method (Kay et al., 2016) implemented in `ripple_detection` (github.com/Eden-Kramer-Lab/ripple_detection). The algorithm was run with a speed threshold of 4 cm s*^−^*^1^, z-score threshold of 2, smoothing *σ* =4 ms, and minimum ripple duration of 15 ms.

**Ripple rates** were calculated on a per-epoch basis as the number of ripple events during the optogenetic test and control intervals divided by the duration of stationary times in the corresponding interval.

##### Sharp Wave Ripple Content

As sharp wave ripples frequently decode to non-local positions, we cannot simply filter times when the animal’s physical location is in a region of the track with sufficient place field coverage. For this analysis, we exclude epochs where at least half of the timepoints cannot be reliably decoded.

A ripple event was classified as non-local if the average decode to animal distance during the interval was greater than 20 cm. Each of these non-local ripples was further classified as continuous/fragmented if the decoding transition state probability was *>*80% weight toward the corresponding state for at least 80% of the ripple duration. It was otherwise labeled as a ‘mixed’ ripple event.

### Data and code availability

All raw data, intermediate analysis files, and code are publicly available to support full reproducibility. Electrophysiology and behavioral data are archived in Neurodata Without Borders (NWB) format (Rübel et al., 2022) via the DANDI Archive (dandiarchive.org/dandiset/001634). Code repositories developed in this work are cited throughout the corresponding methods sections throughout. In preparing this dataset, we developed an NWB extension to store metadata for optogenetics experiments, such as virus, fiber, and stimulation protocol details. This extension is designed for public use and is available at https://github.com/rly/ndx-optogenetics. We built upon this extension in our lab’s custom extension (https://github.com/LorenFrankLab/ndx-franklab-novela) to capture additional metadata specific to our lab’s stimulation protocols. In addition, a complete, containerized analysis pipeline is available at https://github.com/LorenFrankLab/ms_stim_analysis, with Docker images for the database (https://hub.docker.com/r/samuelbray32/spyglass-db-ms_stim_analysis) and notebook environment (https://hub.docker.com/r/samuelbray32/spyglass-hub-ms_stim_analysis).

## References

Atherton, L. A., Dupret, D., and Mellor, J. R. (2015). Memory trace replay: The shaping of memory consolidation by neuromodulation. Trends in Neurosciences, 38(9):560–570.

Bender, F., Gorbati, M., Cadavieco, M. C., Denisova, N., Gao, X., Holman, C., Korotkova, T., and Ponomarenko, A. (2015). Theta oscillations regulate the speed of locomotion via a hippocampus to lateral septum pathway. Nature Communications, 6:8521.

Berg, R. W., Whitmer, D., and Kleinfeld, D. (2006). Exploratory whisking by rat is not phase locked to the hippocampal theta rhythm. The Journal of Neuroscience, 26(24):6518–6522.

Brandon, M. P., Koenig, J., Leutgeb, J. K., and Leutgeb, S. (2014). New and distinct hippocampal place codes are generated in a new environment during septal inactivation. Neuron, 82(4):789–796.

Buño, W. and Velluti, J. C. (1977). Relationships of hippocampal theta cycles with bar pressing during self-stimulation. Physiology & Behavior, 19(5):615–621.

Burgess, N., Recce, M., and O’Keefe, J. (1994). A model of hippocampal function. Neural Networks, 7(6–7):1065–1081.

Buzsáki, G. (1989). Two-stage model of memory trace formation: A role for “noisy” brain states. Neuroscience, 31(3):551–570.

Buzsáki, G. (2002). Theta oscillations in the hippocampus. Neuron, 33(3):325–340.

Buzsáki, G., Leung, L. W., and Vanderwolf, C. H. (1983). Cellular bases of hippocampal EEG in the behaving rat. Brain Research Reviews, 6(2–3):139–171.

Comrie, A. E., Frank, L. M., and Kay, K. (2022). Imagination as a fundamental function of the hippocampus. Philosophical Transactions of the Royal Society B: Biological Sciences, 377(1866):20210336.

Comrie, A. E., Monroe, E. J., Kahn, A. E., Denovellis, E. L., Joshi, A., Guidera, J. A., Krausz, T. A., Berke, J. D., Daw, N. D., and Frank, L. M. (2024). Hippocampal representations of alternative possibilities are flexibly generated to meet cognitive demands. bioRxiv.

Davidson, T. J., Kloosterman, F., and Wilson, M. A. (2009). Hippocampal replay of extended experience. Neuron, 63(4):497–507.

Denovellis, E. L., Gillespie, A. K., Coulter, M. E., Sosa, M., Chung, J. E., Eden, U. T., and Frank, L. M. (2021). Hippocampal replay of experience at real-world speeds. eLife, 10:e64505.

Diba, K. and Buzsáki, G. (2007). Forward and reverse hippocampal place-cell sequences during ripples. Nature Neuroscience, 10(10):1241–1242.

Donahue, M. M., Robson, E., and Colgin, L. L. (2025). Hippocampal place cell sequences are impaired in a rat model of Fragile X syndrome. Journal of Neuroscience, 45(15):e1978242025.

Dragoi, G. and Buzsáki, G. (2006). Temporal encoding of place sequences by hippocampal cell assemblies. Neuron, 50(1):145–157.

Drieu, C., Todorova, R., and Zugaro, M. (2018). Nested sequences of hippocampal assemblies during behavior support subsequent sleep replay. Science, 362(6415):675–679.

Ego-Stengel, V. and Wilson, M. A. (2010). Disruption of ripple-associated hippocampal activity during rest impairs spatial learning in the rat. Hippocampus, 20(1):1–10.

Eichenbaum, H. (2017). Prefrontal-hippocampal interactions in episodic memory. Nature Reviews Neuroscience, 18(9):547–558.

Etter, G., van der Veldt, S., Choi, J., and Williams, S. (2023). Optogenetic frequency scrambling of hippocampal theta oscillations dissociates working memory retrieval from hippocampal spatiotemporal codes. Nature Communications, 14(1):410.

Farrell, J. S., Lovett-Barron, M., Klein, P. M., Sparks, F. T., Gschwind, T. A., Ortiz, A., Ahanonu, B., Terada, S., Bradbury, S., Oijala, M., Hwaun, E., Dudok, B., Szabo, G., Schnitzer, M. J., Deisseroth, K., Losonczy, A., and Soltesz, I. (2021). Supramammillary regulation of locomotion and hippocampal activity. Science, 374(6574):1492–1496.

Fernández-Ruiz, A., Oliva, A., de Oliveira, E. F., Rocha-Almeida, F., Tingley, D., and Buzsáki, G. (2019). Long-duration hippocampal sharp wave ripples improve memory. Science, 364(6445):1082–1086.

Forli, A., Fan, W., Qi, K. K., and Yartsev, M. M. (2025). Replay and representation dynamics in the hippocampus of freely flying bats. Nature, 645(8082):974–980.

Foster, D. J. and Wilson, M. A. (2006). Reverse replay of behavioural sequences in hippocampal place cells during the awake state. Nature, 440(7084):680–683.

Foster, D. J. and Wilson, M. A. (2007). Hippocampal theta sequences. Hippocampus, 17(11):1093–1099.

Freund, T. F. and Antal, M. (1988). GABA-containing neurons in the septum control inhibitory interneurons in the hippocampus. Nature, 336(6195):170–173.

Fuhrmann, F., Justus, D., Sosulina, L., Kaneko, H., Beutel, T., Friedrichs, D., Schoch, S., Schwarz, M. K., Fuhrmann, M., and Remy, S. (2015). Locomotion, theta oscillations, and the speed-correlated firing of hippocampal neurons are controlled by a medial septal glutamatergic circuit. Neuron, 86(5):1253–1264.

Gemzik, Z. M., Donahue, M. M., and Griffin, A. L. (2021). Optogenetic suppression of the medial septum impairs working memory maintenance. Learning & Memory, 28(10):361–370.

Gillespie, A. K., Astudillo Maya, D. A., Denovellis, E. L., Desse, S., and Frank, L. M. (2024). Neurofeedback training can modulate task-relevant memory replay rate in rats. eLife, 12:RP90944.

Gillespie, A. K., Astudillo Maya, D. A., Denovellis, E. L., Liu, D. F., Kastner, D. B., Coulter, M. E., Roumis, D. K., Eden, U. T., and Frank, L. M. (2021). Hippocampal replay reflects specific past experiences rather than a plan for subsequent choice. Neuron, 109(19):3149–3163.e6.

Gupta, A. S., van der Meer, M. A. A., Touretzky, D. S., and Redish, A. D. (2012). Segmentation of spatial experience by hippocampal *θ* sequences. Nature Neuroscience, 15(7):1032–1039.

Hasselmo, M. E. (2025). Development of the SPEAR model: Separate phases of encoding and retrieval are necessary for storing multiple overlapping associative memories. Hippocampus, 35(1):e23676.

Hasselmo, M. E., Bodelón, C., and Wyble, B. P. (2002). A proposed function for hippocampal theta rhythm: Separate phases of encoding and retrieval enhance reversal of prior learning. Neural Computation, 14(4):793–817.

Ito, H. T., Zhang, S.-J., Witter, M. P., Moser, E. I., and Moser, M.-B. (2015). A prefrontal-thalamo-hippocampal circuit for goal-directed spatial navigation. Nature, 522(7554):50–55.

Jadhav, S. P., Kemere, C., German, P. W., and Frank, L. M. (2012). Awake hippocampal sharp-wave ripples support spatial memory. Science, 336(6087):1454–1458.

Jensen, O. and Lisman, J. E. (1996). Theta/gamma networks with slow NMDA channels learn sequences and encode episodic memory: Role of NMDA channels in recall. Learning & Memory, 3(2–3):264–278.

Johnson, A. and Redish, A. D. (2007). Neural ensembles in CA3 transiently encode paths forward of the animal at a decision point. The Journal of Neuroscience, 27(45):12176–12189.

Jones, G. A., Norris, S. K., and Henderson, Z. (1999). Conduction velocities and membrane properties of different classes of rat septohippocampal neurons recorded in vitro. The Journal of Physiology, 517(3):867–877.

Joo, H. R. and Frank, L. M. (2018). The hippocampal sharp wave-ripple in memory retrieval for immediate use and consolidation. Nature Reviews Neuroscience, 19(12):744–757.

Joshi, A. (2017). *Behaviour-Dependent Activity and Synaptic Organisation of Septo-Hippocampal GABAergic Neurons*. PhD thesis, University of Oxford.

Joshi, A., Denovellis, E. L., Mankili, A., Meneksedag, Y., Davidson, T. J., Gillespie, A. K., Guidera, J. A., Roumis, D., and Frank, L. M. (2023). Dynamic synchronization between hippocampal representations and stepping. Nature, 617(7959):125–131.

Joshi, A., Salib, M., Viney, T. J., Dupret, D., and Somogyi, P. (2017). Behavior-dependent activity and synaptic organization of septo-hippocampal GABAergic neurons selectively targeting the hippocampal CA3 area. Neuron, 96(6):1342–1357.e5.

Joshi, A. and Somogyi, P. (2020). Changing phase relationship of the stepping rhythm to neuronal oscillatory theta activity in the septo-hippocampal network of mice. Brain Structure & Function, 225(2):871–879.

Karlsson, M. P. and Frank, L. M. (2009). Awake replay of remote experiences in the hippocampus. Nature Neuroscience, 12(7):913–918.

Kaur, J., Komi, S. A., Dmytriyeva, O., Houser, G. A., Bonfils, M. C. A., and Berg, R. W. (2025). Pedunculopontine-stimulation obstructs hippocampal theta rhythm and halts movement. Scientific Reports, 15(1):17903.

Kay, K., Chung, J. E., Sosa, M., Schor, J. S., Karlsson, M. P., Larkin, M. C., Liu, D. F., and Frank, L. M. (2020). Constant sub-second cycling between representations of possible futures in the hippocampus. Cell, 180(3):552–567.e25.

Kay, K., Sosa, M., Chung, J. E., Karlsson, M. P., Larkin, M. C., and Frank, L. M. (2016). A hippocampal network for spatial coding during immobility and sleep. Nature, 531(7593):185–190.

Kim, J. J. and Fanselow, M. S. (1992). Modality-specific retrograde amnesia of fear. Science, 256(5057):675–677.

Kim, S. M. and Frank, L. M. (2009). Hippocampal lesions impair rapid learning of a continuous spatial alternation task. PLoS ONE, 4(5):e5494.

Király, B., Domonkos, A., Jelitai, M., Lopes-Dos-Santos, V., Martínez-Bellver, S., Kocsis, B., Schlingloff, D., Joshi, A., Salib, M., Fiáth, R., Barthó, P., Ulbert, I., Freund, T. F., Viney, T. J., Dupret, D., Varga, V., and Hangya, B. (2023). The medial septum controls hippocampal supra-theta oscillations. Nature Communications, 14(1):6159.

Kloc, M. L., Velasquez, F., Niedecker, R. W., Barry, J. M., and Holmes, G. L. (2020). Disruption of hippocampal rhythms via optogenetic stimulation during the critical period for memory development impairs spatial cognition. Brain Stimulation, 13(6):1535–1547.

Koenig, J., Linder, A. N., Leutgeb, J. K., and Leutgeb, S. (2011). The spatial periodicity of grid cells is not sustained during reduced theta oscillations. Science, 332(6029):592–595.

Ledberg, A. and Robbe, D. (2011). Locomotion-related oscillatory body movements at 6–12 Hz modulate the activity of hippocampal neurons. PLOS ONE, 6(11):e27575.

Lee, K. H., Denovellis, E. L., Ly, R., Magland, J., Soules, J., Comrie, A. E., Gramling, D. P., Guidera, J. A., Nevers, R., Adenekan, P., Brozdowski, C., Bray, S. R., Monroe, E., Bak, J. H., Coulter, M. E., Sun, X., Broyles, E., Shin, D., Chiang, S., Holobetz, C., Tritt, A., Rübel, O., Nguyen, T., Yatsenko, D., Chu, J., Kemere, C., Garcia, S., Buccino, A., and Frank, L. M. (2024). Spyglass: A framework for reproducible and shareable neuroscience research. bioRxiv preprint.

Lepperød, M. E., Christensen, A. C., Lensjø, K. K., Buccino, A. P., Yu, J., Fyhn, M., and Hafting, T. (2021). Optogenetic pacing of medial septum parvalbumin-positive cells disrupts temporal but not spatial firing in grid cells. Science Advances, 7(19):eabd5684.

Lisman, J. E. and Jensen, O. (2013). The theta-gamma neural code. Neuron, 77(6):1002–1016.

Liu, C., Todorova, R., Tang, W., Oliva, A., and Fernandez-Ruiz, A. (2023). Associative and predictive hippocampal codes support memory-guided behaviors. Science, 382(6668):eadi8237.

Martorell, A. J., Paulson, A. L., Suk, H.-J., Abdurrob, F., Drummond, G. T., Guan, W., Young, J. Z., Kim, D. N.-W., Kritskiy, O., Barker, S. J., Mangena, V., Prince, S. M., Brown, E. N., Chung, K., Boyden, E. S., Singer, A. C., and Tsai, L.-H. (2019). Multi-sensory gamma stimulation ameliorates Alzheimer’s-associated pathology and improves cognition. Cell, 177(2):256–271.e22.

Mizumori, S. J., Perez, G. M., Alvarado, M. C., Barnes, C. A., and McNaughton, B. L. (1990). Reversible inactivation of the medial septum differentially affects two forms of learning in rats. Brain Research, 528(1):12–20.

Mizuseki, K., Sirota, A., Pastalkova, E., and Buzsáki, G. (2009). Theta oscillations provide temporal windows for local circuit computation in the entorhinal-hippocampal loop. Neuron, 64(2):267–280.

Morris, R. G., Garrud, P., Rawlins, J. N., and O’Keefe, J. (1982). Place navigation impaired in rats with hippocampal lesions. Nature, 297(5868):681–683.

Mouchati, P. R., Kloc, M. L., Holmes, G. L., White, S. L., and Barry, J. M. (2020). Optogenetic “low-theta” pacing of the septohippocampal circuit is sufficient for spatial goal finding and is influenced by behavioral state and cognitive demand. Hippocampus, 30(11):1167–1193.

O’Keefe, J. (1976). Place units in the hippocampus of the freely moving rat. Experimental Neurology, 51(1):78–109.

O’Keefe, J. and Recce, M. L. (1993). Phase relationship between hippocampal place units and the EEG theta rhythm. Hippocampus, 3(3):317–330.

Panoz-Brown, D., Iyer, V., Carey, L. M., Sluka, C. M., Rajic, G., Kestenman, J., Gentry, M., Brotheridge, S., Somekh, I., Corbin, H. E., Tucker, K. G., Almeida, B., Hex, S. B., Garcia, K. D., Hohmann, A. G., and Crystal, J. D. (2018). Replay of episodic memories in the rat. Current Biology, 28(10):1628–1634.e7.

Pastalkova, E., Itskov, V., Amarasingham, A., and Buzsáki, G. (2008). Internally generated cell assembly sequences in the rat hippocampus. Science, 321(5894):1322–1327.

Petersen, P. C. and Buzsáki, G. (2020). Cooling of medial septum reveals theta phase lag coordination of hippocampal cell assemblies. Neuron, 107(4):731–744.e3.

Petsche, H., Stumpf, C., and Gogolak, G. (1962). The significance of the rabbit’s septum as a relay station between the midbrain and the hippocampus. Electroencephalography and Clinical Neurophysiology, 14:202–211.

Pfeiffer, B. E. and Foster, D. J. (2013). Hippocampal place-cell sequences depict future paths to remembered goals. Nature, 497(7447):74–79.

Quirk, C. R., Zutshi, I., Srikanth, S., Fu, M. L., Marciano, N. D., Wright, M. K., Parsey, D. F., Liu, S., Siretskiy, R. E., Huynh, T. L., Leutgeb, J. K., and Leutgeb, S. (2021). Precisely timed theta oscillations are selectively required during the encoding phase of memory. Nature Neuroscience, 24(11):1614–1627.

Robbe, D. and Buzsáki, G. (2009). Alteration of theta timescale dynamics of hippocampal place cells by a cannabinoid is associated with a loss of hippocampal-prefrontal oscillatory synchrony. The Journal of Neuroscience, 29(40):12597–12605.

Robinson, J., Manseau, F., Ducharme, G., Amilhon, B., Vigneault, E., El Mestikawy, S., and Williams, S. (2016). Optogenetic activation of septal glutamatergic neurons drive hippocampal theta rhythms. The Journal of Neuroscience, 36(10):3016–3023.

Robinson, J. C., Wilmot, J. H., and Hasselmo, M. E. (2023). Septo-hippocampal dynamics and the encoding of space and time. Trends in Neurosciences, 46(9):712–725.

Robinson, J. C., Ying, J., Hasselmo, M. E., and Brandon, M. P. (2024). Optogenetic silencing of medial septal GABAergic neurons disrupts grid cell spatial and temporal coding in the medial entorhinal cortex. Cell Reports, 43(8):114529.

Rübel, O., Tritt, A., Ly, R., Dichter, B. K., Ghosh, S., Niu, L., Baker, P., Soltesz, I., Ng, L., Svoboda, K., Frank, L., and Bouchard, K. E. (2022). The neurodata without borders ecosystem for neurophysiological data science. eLife, 11:e78362.

Salib, M., Joshi, A., Katona, L., Howarth, M., Micklem, B. R., Somogyi, P., and Viney, T. J. (2019). GABAergic medial septal neurons with low-rhythmic firing innervating the dentate gyrus and hippocampal area CA3. The Journal of Neuroscience, 39(23):4527–4549.

Scoville, W. B. and Milner, B. (1957). Loss of recent memory after bilateral hippocampal lesions. *Journal of Neurology*, Neurosurgery, and Psychiatry, 20(1):11–21.

Siegle, J. H. and Wilson, M. A. (2014). Enhancement of encoding and retrieval functions through theta phase-specific manipulation of hippocampus. eLife, 3:e03061.

Skaggs, W. E., McNaughton, B. L., Wilson, M. A., and Barnes, C. A. (1996). Theta phase precession in hippocampal neuronal populations and the compression of temporal sequences. Hippocampus, 6(2):149–172.

Smith, A. C., Frank, L. M., Wirth, S., Yanike, M., Hu, D., Kubota, Y., Graybiel, A. M., Suzuki, W. A., and Brown, E. N. (2004). Dynamic analysis of learning in behavioral experiments. The Journal of Neuroscience, 24(2):447–461.

Tang, W., Mei, X., Harvey, R. E., Carbajal-Leon, E., Netzer, T., Chang, H., Oliva, A., and Fernandez-Ruiz, A. (2026). Goal-directed hippocampal theta sweeps during memory-guided navigation. Nature Neuroscience, 29(9):2214–2224.

Vancura, B., Geiller, T., Grosmark, A., Zhao, V., and Losonczy, A. (2023). Inhibitory control of sharp-wave ripple duration during learning in hippocampal recurrent networks. Nature Neuroscience, 26(5):788–797.

Varga, C., Golshani, P., and Soltesz, I. (2012). Frequency-invariant temporal ordering of interneuronal discharges during hippocampal oscillations in awake mice. Proceedings of the National Academy of Sciences, 109(40):E2726–E2734.

Viana da Silva, S., Haberl, M. G., Gaur, K., Patel, R., Narayan, G., Ledakis, M., Fu, M. L., de Castro Vieira, M., Koo, E. H., Leutgeb, J. K., and Leutgeb, S. (2024). Localized APP expression results in progressive network dysfunction by disorganizing spike timing. Neuron, 112(1):124–140.e6.

Viney, T. J., Lasztoczi, B., Katona, L., Crump, M. G., Tukker, J. J., Klausberger, T., and Somogyi, P. (2013). Network state-dependent inhibition of identified hippocampal CA3 axo-axonic cells in vivo. Nature Neuroscience, 16(12):1802–1811.

Viney, T. J., Salib, M., Joshi, A., Unal, G., Berry, N., and Somogyi, P. (2018). Shared rhythmic subcortical GABAergic input to the entorhinal cortex and presubiculum. eLife, 7:e34395.

Vollan, A. Z., Gardner, R. J., Moser, M.-B., and Moser, E. I. (2025). Left–right-alternating theta sweeps in entorhinal–hippocampal maps of space. Nature, 639(8056):995–1005.

Wang, Y., Romani, S., Lustig, B., Leonardo, A., and Pastalkova, E. (2015). Theta sequences are essential for internally generated hippocampal firing fields. Nature Neuroscience, 18(2):282–288.

Wang, Y., Roth, Z., and Pastalkova, E. (2016). Synchronized excitability in a network enables generation of internal neuronal sequences. eLife, 5:e20697.

Wikenheiser, A. M. and Redish, A. D. (2015). Hippocampal theta sequences reflect current goals. Nature Neuroscience, 18(2):289–294.

Yong, H. C., Chang, H., and Brandon, M. P. (2022). Optogenetic reduction of theta oscillations reveals that a single reliable time cell sequence is not required for working memory. bioRxiv preprint.

Yu, C., Ji, Z., Ormond, J., O’Keefe, J., and Burgess, N. (2026). Hippocampal theta sweeps indicate goal direction during navigation. Nature Neuroscience, 29(9):2225–2236.

Yu, J. Y., Pettibone, J. R., Guo, C., Zhang, S., Saunders, T. L., Hughes, E. D., Filipiak, W. E., Zeidler, M. G., Bender, K. J., Hopf, F., Smyth, C. N., Kharazia, V., Kiseleva, A., Davidson, T. J., Frank, L. M., and Berke, J. D. (2018). Knock-in rats expressing Cre and Flp recombinases at the parvalbumin locus. bioRxiv preprint.

Zutshi, I., Brandon, M. P., Fu, M. L., Donegan, M. L., Leutgeb, J. K., and Leutgeb, S. (2018). Hippocampal neural circuits respond to optogenetic pacing of theta frequencies by generating accelerated oscillation frequencies. Current Biology, 28(8):1179–1188.e3.

